# α2A-AR–Kv dysfunction drives LC hyperactivity and early sleep disturbance in amyloidogenic mice

**DOI:** 10.1101/2025.09.26.678734

**Authors:** Yi-Ci Zhang, Xue-Ting Zhang, Peng-Yue Chen, Zi-Yue Zhou, Mao-Qing Huang, Kai-Wen He

**Affiliations:** Interdisciplinary Research Center on Biology and Chemistry, Shanghai Institute of Organic Chemistry, Chinese Academy of Sciences, 100 Haike Rd. Shanghai 201203, China; Shanghai Key Laboratory of Aging Studies, Shanghai 201210, China; University of Chinese Academy of Sciences, Beijing 100049, China

**Keywords:** Sleep, Locus coeruleus, Hyperexcitability, Kv4, Kv7, α2A adrenergic receptor, Retigabine

## Abstract

Disrupted sleep–wake patterns are common in neurodegenerative disorders such as Alzheimer’s disease (AD), can emerge early, and are proposed as potent risk factors for disease onset and progression. However, the underlying mechanisms remain poorly understood. Here, we report that 5xFAD transgenic mice exhibit hyperarousal and reduced brain-state transitions, particularly during the dark phase, as early as two months of age. The Locus Coeruleus (LC), a key regulator of arousal and brain-state transitions and a region highly vulnerable in AD, shows time-specific hyperactivity during this phase. This increased tonic LC activity is mediated by heightened neuronal excitability due to impaired Kv4 and Kv7 potassium channel conductance. Pharmacological activation of α2A adrenergic receptors restored Kv4 and Kv7 function and normalized LC activity. Furthermore, local administration of the α2A agonist guanfacine or the Kv7 positive allosteric modulator retigabine substantially rescued the sleep–wake disturbances in young 5xFAD mice. These findings identify dark-phase–selective LC hyperexcitability as a key driver of early-onset sleep disruption in AD mice and implicate α2A adrenergic receptors and Kv7 channels as promising targets for early intervention.

## INTRODUCTION

Sleep-wake disturbances such as hyperarousal and shortened sleep are commonly observed in patients with Alzheimer’s disease (AD)^1–4^. Notably, mounting evidence from longitudinal studies and meta-analyses suggests that sleep disruption during the preclinical phase of AD is not merely a symptom but may actively contribute to the pathogenesis of the disease. Therefore, early sleep disturbances are increasingly recognized as a modifiable risk factor for AD^1, 3, 5^. However, the underlying mechanisms driving these early alterations in sleep–wake regulation remain poorly understood.

The locus coeruleus (LC), a small pontine nucleus composed predominantly of tyrosine hydroxylase-positive (TH+) noradrenergic neurons, plays a central role in regulating arousal, sleep–wake transitions, and attention^6–9^. Optogenetics activation of LC neurons has been shown to promote wakefulness and suppress non-rapid eye movement, NREM, sleep in a frequency-dependent manner^10^. Beyond its well-established role in initiating arousal, recent studies have highlighted the nuanced, state-dependent dynamics of LC neuronal activity that gate transitions between brain states. Moreover, prolonged activation of LC neurons leads to activity-dependent fatigue mediated by α2A-adrenergic receptor (α2A-AR)–driven auto-inhibition, contributing to the buildup of sleep pressure^11^.

Importantly, the LC is also one of the earliest and most vulnerable brain regions affected in AD. Postmortem and in vivo imaging studies reveal extensive LC degeneration that correlates with both disease severity and cognitive decline in AD patients^12–14^. Notably, pathological changes in the LC can occur decades before the onset of clinical symptoms^15, 16^, raising the possibility that early LC dysfunction contributes to the earliest stages of AD progression, including sleep disturbances. However, how such early LC abnormalities arise—and their precise consequences for behavior—remains largely unexplored.

In this study, we investigated the relationship between early LC dysfunction and sleep-wake disturbances using 2-month-old 5xFAD mice, a widely used transgenic model of AD amyloidosis that exhibits early Aβ accumulation but minimal neurodegeneration or cognitive decline at this age. We found that young 5xFAD mice display marked hyperarousal and reduced brain state transitions particularly during the dark phase. Electrophysiological analyses revealed a dark phase–selective hyperexcitability of LC neurons, driven by impaired α2A-AR-mediated modulation of Kv4 and Kv7 potassium channel conductance. Pharmacological enhancement of either α2A-AR or Kv7 activity locally in the LC was sufficient to restore normal sleep–wake patterns in these mice.

Together, our findings identify a previously unrecognized mechanism linking early LC hyperactivity to disrupted sleep architecture in the preclinical phase of AD. These results underscore the critical role of noradrenergic dysfunction in early disease-related behavioral phenotypes and suggest that targeting LC excitability may offer a novel therapeutic strategy for early intervention in Alzheimer’s disease.

### RESULT

### Early-onset sleep deficits in 5xFAD mice characterized by hyperarousal and reduced brain state transitions

Sleep disturbances are well-documented in aged mouse models AD^17–19^. Consistent with these findings, we observed a significant reduction in both non-rapid eye movement (NREM) and rapid eye movement (REM) sleep, along with increased wakefulness, in 12-month-old 5xFAD mice (Fig S1A-F). These results, in line with prior studies, underscore the utility of transgenic AD models for studying sleep dysfunctions commonly associated with the disease^3, 20, 21^.

To explore the early sleep abnormalities, we examined 5xFAD mice at two months of age—an early disease stage marked by sparse amyloid deposition (Fig S1N-O), but without reported cognitive deficits^22^. At this stage, the sleep-wake architecture remained relatively normal and intact (Fig 1A-C). However, compared to wild-type (WT) littermates, 5xFAD mice exhibited prolong wakefulness and decreased NREM sleep duration compared to their wild-type littermates, particularly during the dark phase (Fig 1D-F). This was accompanied by a decreased number of wake, NREM, and REM bouts (Fig 1J, L), as well as fewer transitions between wake–NREM and NREM–REM states (Fig S1M) in the dark cycle, suggesting a state of hyperarousal and impaired flexibility in sleep–wake regulation.

**Figure 1.**
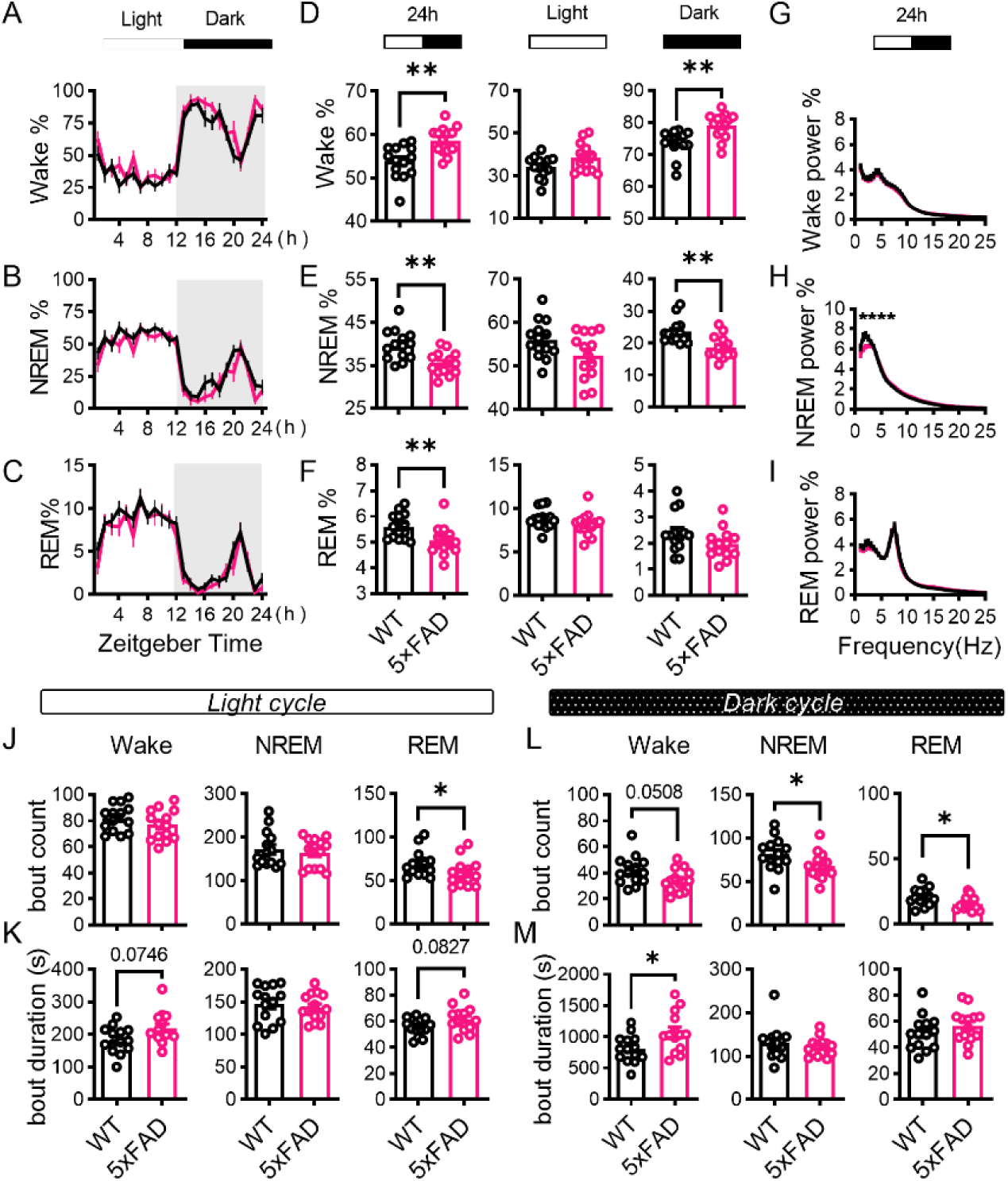
Early-onset sleep deficits in 2-month-old 5xFAD mice. A-F. Percentages of time spent in wake (A), NREM (B), and REM (C) across the light/dark cycle in 2-month-old 5xFAD mice and their WT littermates. Comparisons of the percentage of time spent in wake (D), NREM (E), and REM (F) during the 24-hour day (left panels), light cycle (middle panels) and the dark cycle (right panels) between 2-month-old 5xFAD mice and their WT littermates. (WT: n = 14; 5xFAD: n = 14) G-I. Comparison of the normalized EEG power spectrums of wake (G), NREM (H), and REM (I) during the 24-hour day between 2-month-old 5xFAD mice and WT mice. (WT: n = 14; 5xFAD: n = 14) J-M. The differences in bout count number (J,L) and mean bout duration (K,M) for distinct vigilance stages during light cycle and dark cycle when comparing the 2-month-old 5×FAD mice to their WT controls. Left panels, wake. Middle panels, NREM. Right panels, REM. (WT: n = 14; 5xFAD: n = 14)

Furthermore, young 5xFAD mice showed reduced NREM delta (δ, 0.5–4 Hz) power (Fig 1G–I, Fig S1G), a phenotype observed in both AD patients and aged AD mouse models^23^, while maintaining relatively preserved sleep homeostasis (Fig S1H–K). Collectively, these findings demonstrate the emergence of sleep deficits as early as two months of age in 5xFAD mice, characterized by dark phase-selective hyperarousal, impaired state transitions, and decreased delta power.

### Dark phase-selective hyperactivity of LC neurons in young 5xFAD mice

To investigate the potential cause of early-onset sleep disturbances in young 5xFAD mice, we focused on the locus coeruleus (LC), a brainstem nucleus critical for sleep–wake regulation^7^. Activation of LC neurons promotes arousal^10^. Their activities are recently found to be also important for brain state transitions^24^. Moreover, LC is among the earliest brain regions affected in Alzheimer’s disease (AD)^16^. Therefore, given the observed sleep phenotypes in young 5xFAD mice, we sought to examine LC neuron dynamics in young 5xFAD mice.

LC neuron acts as pacemakers with low tonic firing rates that vary depending on brain states^8^. To assess their activity, we recorded tonic firing rates in acute brain slices collected during the dark phase (zeitgeber time [ZT] 22) and the light phase (ZT9) from 2-month-old 5xFAD mice and wild-type (WT) littermates. LC neurons were identified based on their location and morphology, with their identity confirmed via post-hoc staining for tyrosine hydroxylase (TH) and biotin filling (Fig 2A).

**Figure 2.**
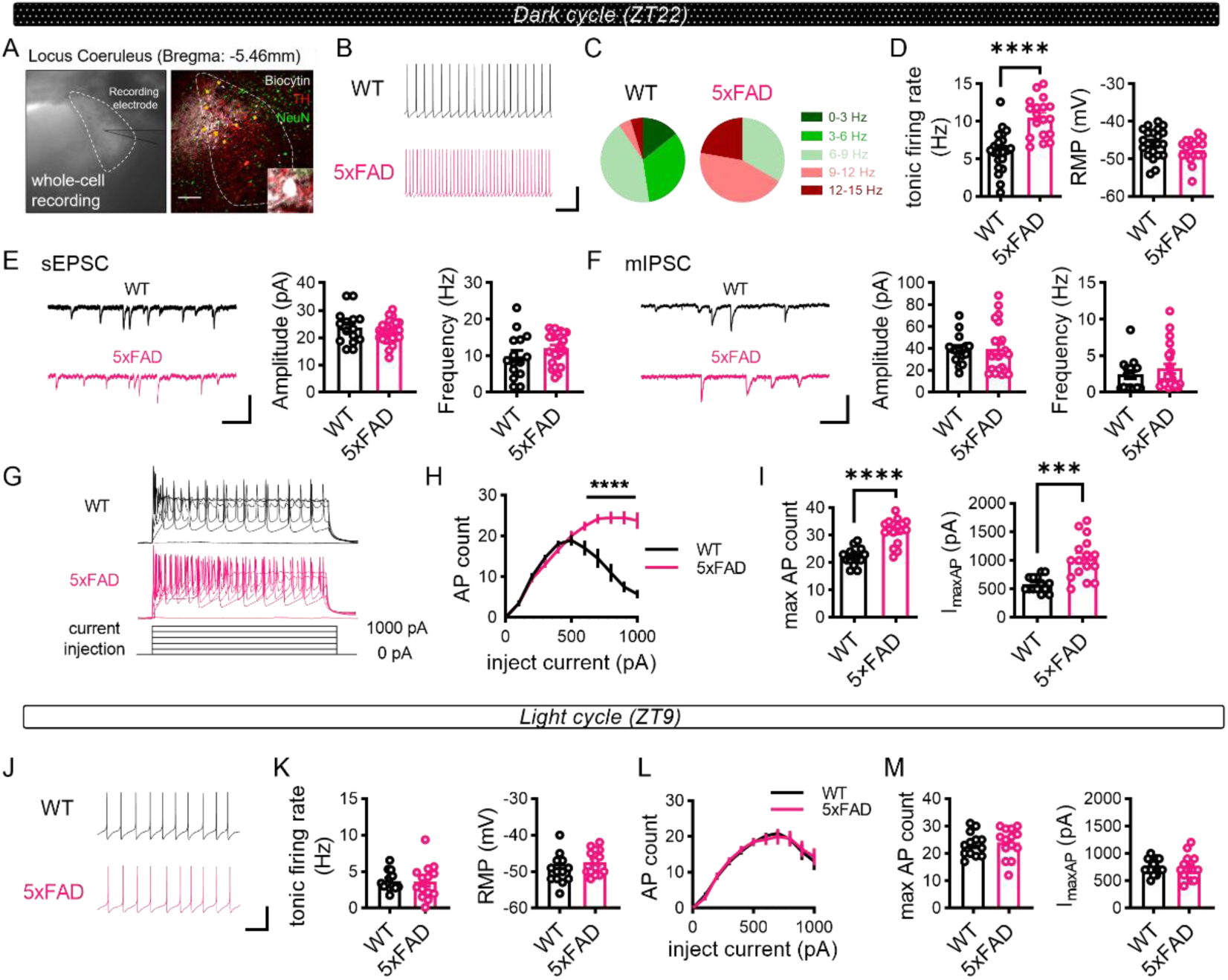
Dark phase-selective hyperactivity and hyperexcitability of LC neurons in 2-month-old 5xFAD mice. A-I. Whole cell recording of LC neurons from acute brain slices prepared at ZT22. A-D. *Ex vivo* recordings of tonic firing of LC neurons in 2-month-old 5xFAD mice and WT littermates (WT: 21 cells from 3 mice; 5xFAD: 18 cells from 3 mice). A, Left, DIC image taken during recording. Right, biocytin-filled recorded cells (arrowhead) co-stained with TH and NeuN. (biocytin: white; TH: red; NeuN: green. Scale bar, 100 µm). B, Representative traces of recording (Scale bar, 600ms, 25pA). C, Distribution of firing rate. D, Quantification of firing rate (left panel) and resting membrane potential (RMP, right panel). E. Comparison of spontaneous EPSCs between 5xFAD and WT (WT: 14 cells from 3 mice; 5xFAD: 16 cells from 3 mice). Left panel, example sEPSC traces. Middle and right panels, quantification of the sEPSC amplitude (Middle) and frequency (Right) (Scale bars, 50pA, 250ms) F. Comparison of miniature IPSCs between 5xFAD and WT (WT: 16 cells from 3 mice; 5xFAD: 22 cells from 3 mice). Left panel, example mIPSC traces. Middle and right panels, quantification of the mIPSC amplitude (Middle) and frequency (Right). (Scale bars, 50pA, 250ms) G-I. Comparison of neuronal excitability by injecting current steps (WT: 14 cells from 3 mice; 5xFAD: 16 cells from 3 mice). G, Representative traces evoked by injecting 500 ms current steps (0, 200, 400, 600, 800, 1000 pA). H, Current-spiking relationship of LC neurons. I, Quantification of maximal firing rate (Left panel) and currents required to generate the maximal firing (I_maxAP_, Right panel). J-M. Whole cell recording of LC neurons from acute brain slices prepared at ZT9. J-K. *Ex vivo* recordings of tonic firing of LC neurons in 2-month-old 5xFAD mice and WT littermates (WT: 13 cells from 3 mice; 5xFAD: 14 cells from 3 mice). J, Representative traces of recording (Scale bar, 600ms, 25pA). K, Quantification of firing rate (left panel) and resting membrane potential (RMP, right panel). L-M. Comparison of neuronal excitability by injecting current steps (WT: 13 cells from 3 mice; 5xFAD: 14 cells from 3 mice). L, Current-spiking relationship of LC neurons. M, Quantification of maximal firing rate (Left panel) and currents required to generate the maximal firing (I_maxAP_, Right panel).

In the dark phase, LC neurons from 5xFAD mice exhibited significantly elevated tonic firing rates compared to WT controls despite the unchanged resting membrane potential (Fig 2B-D) and membrane resistance (Fig S2C). More than 65% of LC neurons in 5xFAD mice fired at rates exceeding 9 Hz compared to less than 10% in wild-type littermates (Fig 2C). This corresponded to a two-fold increase in average tonic firing rate in 5xFAD mice in the dark phase (Fig 2D, left panel). Notably, this hyperactivity was phase-selective: no difference in tonic LC firing rate was observed between genotypes in the light phase (Fig 2J-K). These findings align with the observed dark phase-selective sleep phenotypes and suggest that elevated LC activity may underlie the hyperarousal and impaired transition observed in young 5xFAD mice.

To determine whether the increased LC activity in dark phase stemmed from altered synaptic inputs or neuronal excitability, we first assessed spontaneous excitatory postsynaptic currents (sEPSCs, Fig 2E) and miniature inhibitory postsynaptic currents (mIPSCs, Fig 2F). Neither sEPSCs nor mIPSCs differed between young 5xFAD mice and their wildtype littermates, suggesting unaltered synaptic inputs at this age. We then assessed neuronal excitability. While threshold potentials were similar, LC neurons in 5xFAD mice exhibited slightly reduced rheobase (Fig S2A-B), significantly elevated maximal firing rate (Fig 2I Left panel) and slower spike frequency accommodation in response to more depolarized current steps (Fig 2H). Unlike WT LC neurons, which reached their maximal firing rate with on average 500 pA current injection, many neurons in 5xFAD mice tolerated much higher injected current (I_maxAP_, Fig 2I Right panel).

Importantly, these excitability changes were absent during the light phase (Fig 2L-M), reinforcing the phase-selective nature of LC hyperactivity in young 5xFAD mice. Together, these data suggest that increased intrinsic excitability—rather than altered synaptic input—underlies the dark-phase-specific hyperactivity of LC neurons and may contribute to early sleep–wake disturbances in this AD mouse model.

### Impaired low-threshold Kv4 and Kv7 conductance mediating LC neuronal hyperexcitability

Voltage-gated potassium (Kv) channels play a crucial role in the generation and maintenance of action potential^25^. Therefore, we investigated whether there are alterations in Kv channel conductance in the LC neurons of 5xFAD mice. Kv channels were categorized as either high-threshold (HT) or low-threshold (LT)^25^. We recorded both total and HT K^+^ current (I_k_) by pre-holding at either -120 mV or -30 mV (Fig 3A, see Method). LT K^+^ current was calculated post hoc by subtracting HT from total I_k_ (Fig 3B). Both acute (fast-kinetic, Fig 3B, grey arrow head) and persist (slow-kinetic, Fig 3B, black arrow head) I_k_ amplitudes were quantified. LC neurons from 5xFAD mice showed a significant reduction in total I_k_, including both acute (Fig 3C) and persistent components (Fig 3F), primarily attributed to impaired LT (Fig 3E&H) rather than HT (Fig 3D&G) K^+^ current. Kv4 and Kv7 channels, major contributors to LT acute and persistent currents, respectively^25^, are highly expressed in the LC (Fig S3I). We hypothesized that impaired Kv4 and Kv7 channel properties may underlie the hyperexcitability observed in LC neurons of 5xFAD mice. Pharmacological inhibition of Kv7 with XE991 reduced LT persistent K^+^ conductance (Fig S3A), mimicking the phenotype of 5xFAD neurons and increased neuronal excitability in wild-type mice by elevating the maximal firing rate (Fig 3I). Similarly, inhibition of Kv4 using U0126^26, 27^ suppressed LT acute K^+^ currents (Fig S3B) and increased firing rates by preventing depolarization blockade (Fig 3J). These results suggest that impaired Kv4 and Kv7 channels contribute to the slower accommodation and elevated maximal firing rate in the LC neurons of young 5xFAD mice. Indeed, co-application of Kv4 (NS5806) and Kv7 (retigabine) agonists restored LT I_k_ (Fig S3C-D) and mitigated hyperexcitability (Fig 3K). Activation of Kv4 and Kv7 channels also rescued LC neuron hyperactivity (Fig 3L).

**Figure 3.**
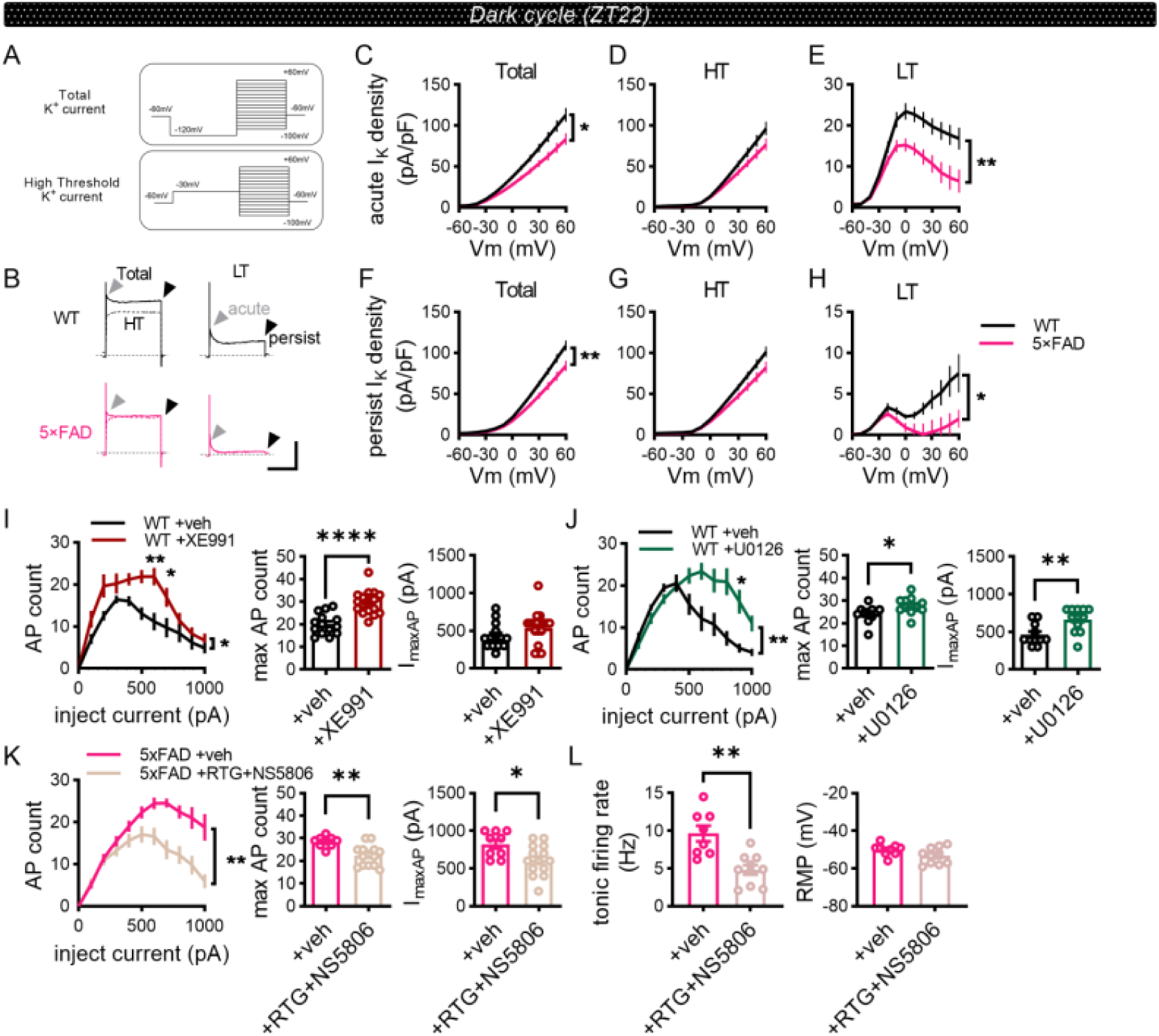
Downregulated Kv4 and Kv7 currents mediate elevated excitability and activity of LC neurons in young 5xFAD mice. A. Diagram depicting protocols used for measuring the total voltage-gated I_k_ (right) and high threshold voltage-gated I_k_ (left). B-H. Comparison of Kv channel conductance of LC neurons between WT and 5xFAD (WT:12 cells from 3 mice; 5xFAD: 13 cells from 3 mice). B, Right panels, Example traces of total (solid traces) and high threshold (HT, dotted traces) potassium current (I_k_). Left panels, Example traces of low threshold (LT) I_k_. Grey arrowheads: acute I_k_. Black arrowheads: persistent I_k_. Scale bar: 2000 pA, 300 ms. C-H, V-I_k_ curves of acute (C,D,E) and persistent (F,G,H) I_k_ density. Left panels, total I_k_; Middle panels, HT I_k_; Right panels, LT I_k_. I-J. Effect of XE991 (I, Veh: 14 cells from 4 mice; XE991: 18 cells from 3 mice) or U0126 (J, Veh: 11 cells from 3 mice; U0126: 12 cells from 3 mice) on neuronal excitability of LC neurons of WT mice. Left panels, current-spiking relationship. Middle panels, maximal firing rate. Right panels, current required to generate the maximal firing (I_maxAP_). K. Effect of co-application of retigabine (RTG) and NS5806 on neuronal excitability of LC neurons of 5xFAD mice (Veh: 9 cells from 3 mice; RTG+NS5806: 14 cells from 3 mice). Left panel, current-spiking relationship. Middle panel, maximal firing rate. Right panel, I_maxAP_. L. Effect of co-application of retigabine (RTG) and NS5806 on tonic firing of LC neurons in 5xFAD mice (Veh: 8 cells from 3 mice; RTG+NS5806: 10 cells from 3 mice). Left panel, tonic firing rate. Right panel, resting membrane potential.

In summary, our findings demonstrate that impaired LT Kv channels, particularly Kv4 and Kv7, mediate the hyperexcitability and hyperactivity of LC neurons in young 5xFAD mice.

### Impaired α2A adrenergic receptor signaling drives LC hyperactivity via dysfunctional Kv4 and Kv7 channel conductance

The impaired Kv4 and Kv7 channel conductance observed in LC neurons of 5xFAD mice may stem from disrupted regulation rather than reduced expression. Bulk RNA sequencing (Fig S3I), qPCR analysis (Fig S3J), and immunofluorescent labeling of major Kv4 and Kv7 subunits (Fig S3K–M) revealed no significant differences in mRNA or total protein levels between 5xFAD and wild-type (WT) mice. These results suggest that the functional impairment arises from altered modulation rather than transcriptional or translational deficits at the early stage.

Given that norepinephrine (NE) is the principal neurotransmitter released by LC neurons, and that NE can exert feedback inhibition via α2-adrenergic autoreceptors located on both synaptic and somatic sites^28, 29^, we explored the role of α2A-adrenergic receptor (α2A-AR) signaling. The α2A-AR subtype is highly expressed in the LC (Fig S4A), and its dysfunction has previously been implicated in promoting LC hyperactivity^30–32^. To assess whether impaired α2A-AR signaling contributes to LC hyperactivity in young 5xFAD mice, we applied guanfacine, a selective α2A-AR agonist, during electrophysiological recordings. Guanfacine significantly reduced tonic firing rates in 5xFAD LC neurons (Fig 4A), but had no effect in WT neurons (Fig 4B), indicating that reduced endogenous α2A-AR activation underlies the increased firing activity observed in 5xFAD mice during the dark phase.

**Figure 4.**
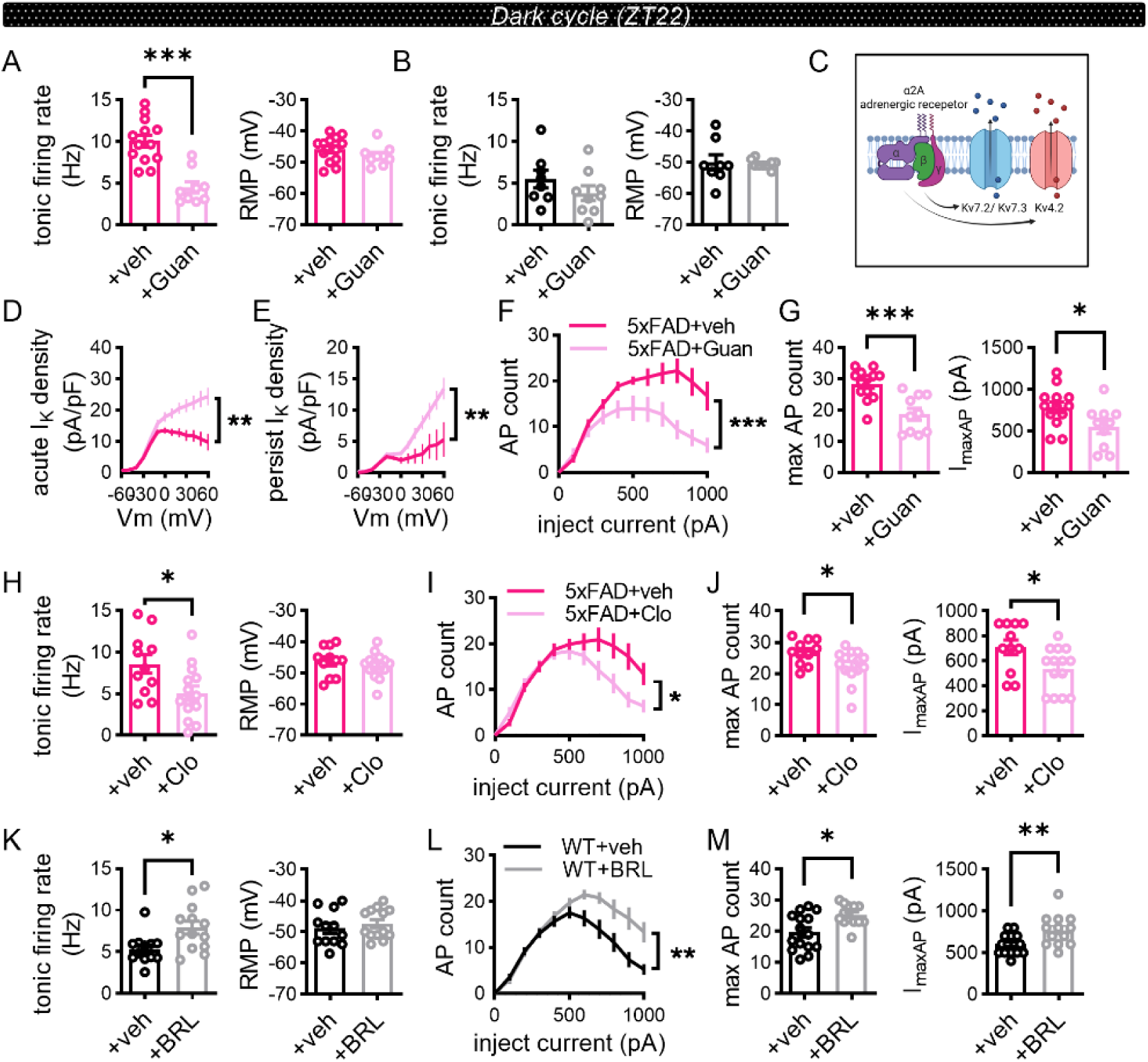
Activation of α2A adrenergic receptor restores Kv channel conductance and LC neuron activity in young 5xFAD mice. A-B. Effect of guanfacine (Guan) on tonic firing of LC neurons in 5xFAD mice (A. Veh: 13 cells from 3 mice; Guan: 9 cells from 3 mice) and WT mice (B. Veh: 8 cells from 3 mice; Guan: 9 cells from 3 mice). Left panel, tonic firing rate. Right panel, resting membrane potential. C. Hypothesized model depicting how α2A adrenergic receptor modulates Kv4 and Kv7 channel properties. D-G. Effect of guanfacine (Guan) on low threshold acute I_k_. (D), low threshold persist I_k_. (E) and neuronal excitability (F-G) of LC neurons in 5xFAD mice (D-E. Veh: 16 cells from 4 mice; Guan: 22 cells from 5 mice; F-G. Veh: 13 cells from 3 mice; Guan: 10 cells from 3 mice) D, low threshold acute I_k_, E, low threshold persist I_k_. F, current-spiking relationship. G, Left panel, maximal firing rate. Right panel, I_maxAP_. H-J. Effect of application of clonidine (Clo) on tonic firing (H) and neuronal excitability (I-J) of LC neurons of 5xFAD mice (Veh: 11 cells from 3 mice; Clo: 14 cells from 3 mice). H, Left panel, tonic firing rate. Right panel, resting membrane potential. I-J, I, current-spiking relationship. J, Left panel, maximal firing rate. Right panel, I_maxAP_. K-M. Effect of BRL-44408 maleate (BRL) on tonic firing (K) and neuronal excitability (L-M) of LC neurons in WT mice (Veh: 15 cells from 3 mice; BRL: 13 cells from 3 mice). K, Left panel, tonic firing rate. Right panel, resting membrane potential. L-M, L, current-spiking relationship. M, Left panel, maximal firing rate. Right panel, I_maxAP_.

Because α2-adrenergic receptors have been shown to modulate Kv7 channel activity ^33–35^, we hypothesized that impaired α2A-AR signaling might mediate the observed reductions in Kv4 and Kv7 conductance (Fig 4C). Indeed, guanfacine application significantly increased both the low-threshold acute and persist I_k_ in LC neurons from 5xFAD mice (Fig 4D-E), restoring normal excitability (Fig 4F-G). These effects were replicated using clonidine, another α2-AR agonist, which similarly reduced neuronal hyperexcitability in 5xFAD mice (Fig 4H-J). Conversely, pharmacological blockade of α2A-ARs using BRL-44408 maleate in WT mice induced a hyperexcitable phenotype resembling that of 5xFAD neurons (Fig 4K-M), while neither clonidine (Fig S4D-F) nor guanfacine (Fig S4G-H) altered WT LC neuronal properties.

Together, these results demonstrate that impaired α2A-AR signaling in young 5xFAD mice leads to decreased Kv4 and Kv7 conductance, resulting in LC hyperexcitability and hyperactivity in the dark phase. This mechanism likely contributes to the early-onset sleep disturbances observed in this AD model.

### Restoration of sleep–wake disturbances in 5xFAD mice via local modulation of LC α2A-AR and Kv channel activity

Our previous findings demonstrated that impaired α2A-AR signaling and reduced downstream Kv4 and Kv7 channel conductance contribute to LC hyperactivity in young 5xFAD mice. To determine whether this signaling dysfunction underlies the early-onset sleep-wake abnormalities observed in this AD model, we employed pharmacological interventions targeting this pathway.

Guanfacine, a clinically approved α2A-AR agonist used to treat various neurological disorders^36, 37^, was locally infused bilaterally into the LC of young 5xFAD mice via implanted cannulas immediately prior to the onset of the dark phase (Fig 5A; see Methods). Compared to vehicle controls, guanfacine significantly reduced wakefulness and increased NREM sleep during the dark phase (Fig 5B-G). In addition, guanfacine-treated mice exhibited a trend toward enhanced transitions between vigilance states (Fig S6A-C), suggesting improved sleep–wake flexibility. These results implicate LC α2A-AR signaling as a key regulator of arousal and sleep architecture, and its dysfunction as a contributor to early-onset sleep disturbances in 5xFAD mice.

**Figure 5.**
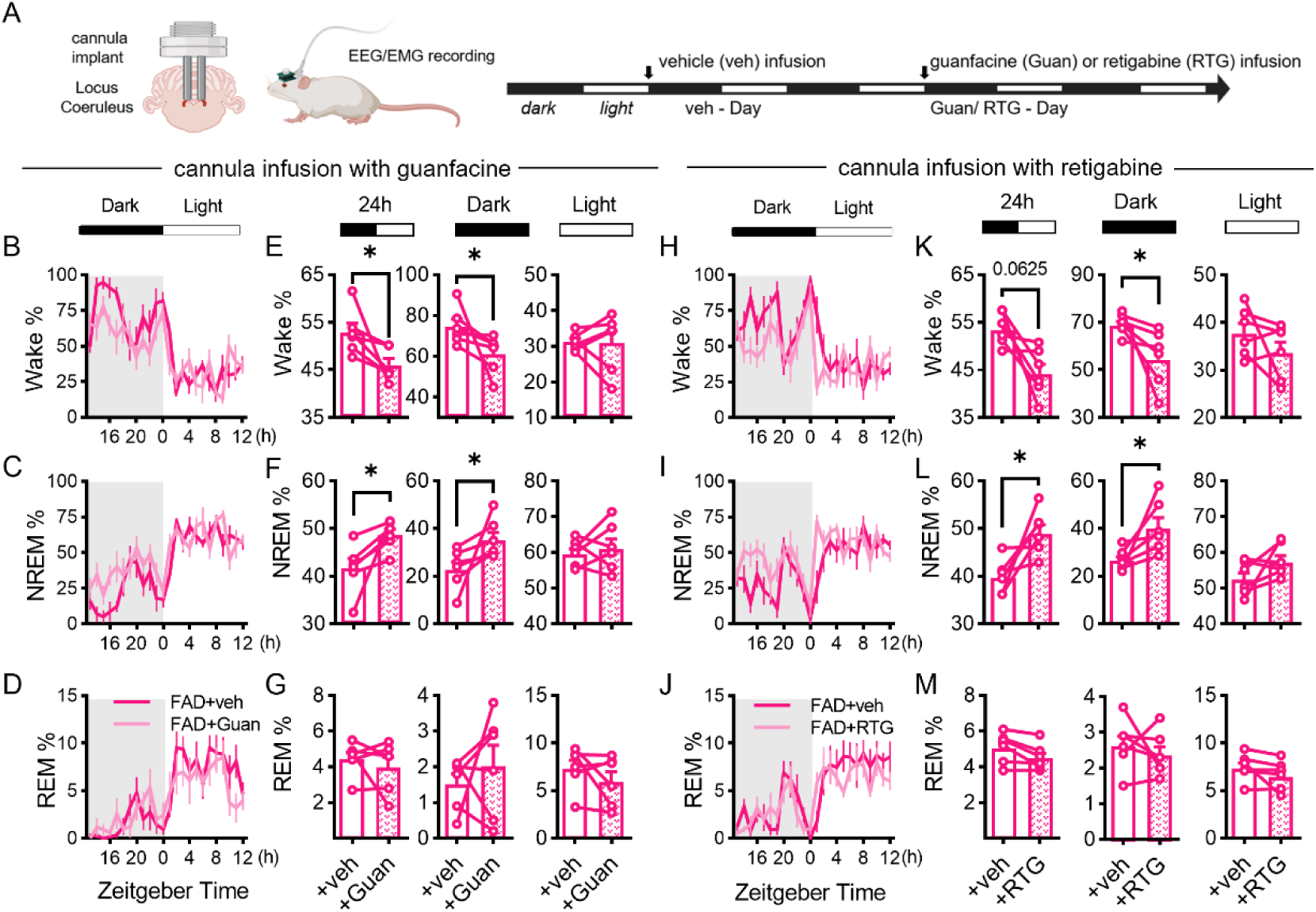
Local infusion of guanfacine or retigabine alleviates the sleep-wake disturbances in 2-month-old 5xFAD mice. A. Schematic diagram showing the experimental design of cannula implantation in LC and the EEG/EMG recording timeline to evaluate the effect of guanfacine or retigabine on sleep pattern. B-G. Comparisons of the percentages of time spent in wake (upper panels), NREM (middle panels), and REM (lower panels) immediately after drugs (vehicle: veh, opened bars; guanfacine: Guan, patterned bars) infusion through cannula at ZT12 (Arrow) in 5xFAD mice. B-D, Percentages of time spent in wake (B), NREM (C), and REM (D) across the light/dark cycle. E-G, Percentages in the 24-hour day (left panels), the dark cycle (middle panels) and the light cycle (right panels) were analyzed. (5xFAD: n=6) H-M. Comparisons of the percentages of time spent in wake (upper panels), NREM (middle panels), and REM (lower panels) immediately after drugs (vehicle: veh, opened bars; retigabine: RTG, patterned bars) infusion into LC areas of 5xFAD mice at ZT12. H-J, Percentages of time spent in wake (H), NREM (I), and REM (J) across the light/dark cycle. K-M, Percentages in the 24-hour day (left panels), the dark cycle (middle panels) and the light cycle (right panels) were analyzed. (5xFAD: n=6)

We next assessed whether direct activation of Kv channels could similarly restore normal sleep architecture. Retigabine, a Kv7 channel opener, was previously shown to reduce LC hyperexcitability ex vivo (Fig S3E-H). Local infusion of retigabine into the LC of 5xFAD mice during the dark phase significantly ameliorated hyperarousal and restored NREM sleep (Fig 5H-M), mirroring the effects of guanfacine. These findings were further confirmed through systemic administration: a single intraperitoneal (IP) injection of retigabine in 5xFAD mice significantly increased NREM sleep and the number of wake–NREM transitions during the dark phase (Fig S7A–M), without altering NREM δ power (Fig S7H–J). Notably, the same treatment had only modest effects on sleep parameters in wild-type mice (Fig S8), suggesting a disease-specific therapeutic potential.

Together, these results support a mechanistic model in which LC hyperactivity—driven by disrupted α2A-AR signaling and impaired Kv channel function—plays a central role in the early manifestation of sleep–wake disturbances in AD. Pharmacological modulation of this pathway may represent a promising strategy for early intervention in AD-associated sleep pathology.

## DISCUSSION

Sleep-wake disturbances are increasingly recognized as an early and clinically significant feature of AD, often preceding the onset of cognitive decline by years. Hyperarousal, sleep fragmentation, and reduced total sleep time are frequently reported in both patients with early-stage AD and individuals at risk. These alterations are not merely epiphenomena of neurodegeneration but may actively contribute to the pathogenesis of AD by exacerbating amyloid-beta (Aβ) accumulation and tau pathology through disrupted glymphatic clearance and elevated neuronal activity^1, 38–40^. Thus, understanding the cellular and circuit-level mechanisms underlying early-onset sleep-wake dysregulation in AD is of both fundamental and translational importance.

Our results identify a key role for dysfunctional α2A-AR-mediated auto-inhibition via modulating Kv 4 and 7 channel conductance in the hyperexcitability of LC neurons in young 5xFAD mice, particularly during the dark (wake-dominant) phase. Under normal physiological conditions, α2A-ARs act as presynaptic^41^ and somatodendritic^42^ autoreceptors that are activated by elevated extracellular norepinephrine (NE) levels, typically during periods of sustained LC activity. These receptors are Gi/o-coupled G protein–coupled receptors (GPCRs) that inhibit further neuronal firing and NE release. However, NE release from somatic sites occurs predominantly at high LC firing rates (>15 Hz), whereas low-frequency activity (<4 Hz) produces sparse somatic NE release^42, 43^. This supports the model in which somatic α2A-ARs primarily function during arouse-dominant dark period when LC neurons are more active (Fig 2D, K)^6, 8^. In line with this model, our data show that in 5xFAD mice, the impaired α2A-AR function leads to pronounced hyperactivity of LC neurons only during the dark phase. In contrast, the light phase, characterized by lower overall LC activity and NE release, remains largely unaffected, masking the defect in α2A-AR signaling during this period.

Although significant reductions in α2A-AR mRNA and its binding sites have been reported in the LC of AD patients at later stages^44^, our data suggest that functional impairment of α2A-AR signaling precedes these transcriptional and translational changes (Fig S4B & C). At this early stage, the deficit may be due to post-translational mechanisms, such as abnormal receptor phosphorylation^45^, membrane trafficking, or downregulation of interacting proteins^46^. Low levels of Aβ accumulation and early-stage neuroinflammation (Fig S5) may contribute to this dysfunction by altering receptor localization or stability^47, 48^. Notably, Aβ oligomers have been shown to bind to an allosteric site on α2-adrenergic receptors, thereby redirecting downstream signaling pathways in a pathological manner^49^. This direct interaction may further disrupt α2A-AR function independent of gene expression levels.

Mechanistically, we found that α2A-AR activation enhances the conductance of both Kv4 and Kv7 channels, and this enhancement is sufficient to suppress the hyperexcitability of LC neurons in young 5xFAD mice. These findings point to the importance of intact α2A-AR–Kv channel coupling in maintaining LC excitability and proper sleep-wake architecture. As a GPCR, α2A-AR activation promotes the dissociation of Gβγ subunits^50^, which has been shown to directly bind to and increase the open probability of Kv7 channel^34^, thereby dampening neuronal excitability.

Importantly, Kv dysfunction is commonly observed in aging and diseases^51, 52^. Both Kv4 and Kv7 channels have been implicated in pathological neuronal hyperexcitability and seizures when their function is compromised^53–55^. Notably, reduced Kv7 activity in hypothalamic hypocretin (Hcrt) neurons has been shown to promote hyperexcitability and sleep disruption in aged mice^56^, supporting a broader role for Kv dysfunction in sleep and arousal disturbances. Beyond direct α2A-AR signaling deficits, other mechanisms may contribute to the impaired Kv channel function observed in early AD model. These include post-translational modifications that alter channel kinetics^57, 58^, downregulation of auxiliary or interacting proteins necessary for channel function^59^, disrupted subcellular trafficking and localization of Kv channels^60^, and changes of micro-environment and signaling pathways that modulate channel expression or activity^61^. With more drugs targeting Kv channels become available^62^, in-depth analysis of which and how Kv channels are dysregulated in early AD is necessary.

The sleep-wake disturbances observed in 5xFAD mice—hyperarousal, inflexible state transitions, and prolonged wake bouts—mirror clinical observations in individuals with mild cognitive impairment (MCI) and prodromal AD. In human studies, early-stage AD patients frequently exhibit elevated arousal levels, increased sleep latency, and heightened anxiety symptoms^63, 64^. LC hyperactivity may be a converging mechanism underlying these phenotypes, as the LC mediates arousal, stress reactivity, and anxiety via widespread NE projections to the cortex, hippocampus, and amygdala^8, 65^. More importantly, the initial hyperactivity of LC neurons may have long-term consequences for AD progression^63, 64^. Recent finding indicates that the hyperactivity of LC neurons in APP knock-in mice leads to the early degradation of LC fibers in the olfactory bulk that drives the olfactory dysfunction^66^. These observations position early LC hyperactivity as both a biomarker and a potential driver of AD progression.

Taken together, our findings highlight the pivotal role of LC dysregulation—specifically α2A-AR signaling deficits and associated Kv channel dysfunction—in mediating early sleep-wake disturbances in AD. Therapeutically, restoring LC auto-inhibition or directly targeting LC hyperexcitability may represent promising early interventions to alleviate arousal dysregulation associated with the prodromal phase.

## ACKNOWLEDGEMENT

We thank the staff members of the animal facility at the National Facility for Protein Science in Shanghai (NFPS), Shanghai Advanced Research Institute, Chinese Academy of Sciences, China for excellent support. This work is supported by Shanghai Science and Technology Development Funds (Grant No. 22ZR1475100), NSFC (Grant No. 32070963, 32271004), and Shanghai Municipal Science and Technology Major Project (Grant No. 2019SHZDZX02) to K.-W.H.

## AUTHOR CONTRIBUTIONS

Conceptualization, K.-W.H.; Primary Investigation, Y.-C. Z.; Assistant Investigation, X.-T. Z., P.-Y. C., Z.-Y. Z., M.-Q. H.; Data Analysis, Y.-C. Z., X.-T. Z., Z.-Y. Z.; Writing, Y.-C. Z. and K.-W.H.; Funding Acquisition, K.-W.H; Supervision, K.-W.H.

## SUPPLEMENTARY FIGURES AND LEGENDS

**Figure S1.**
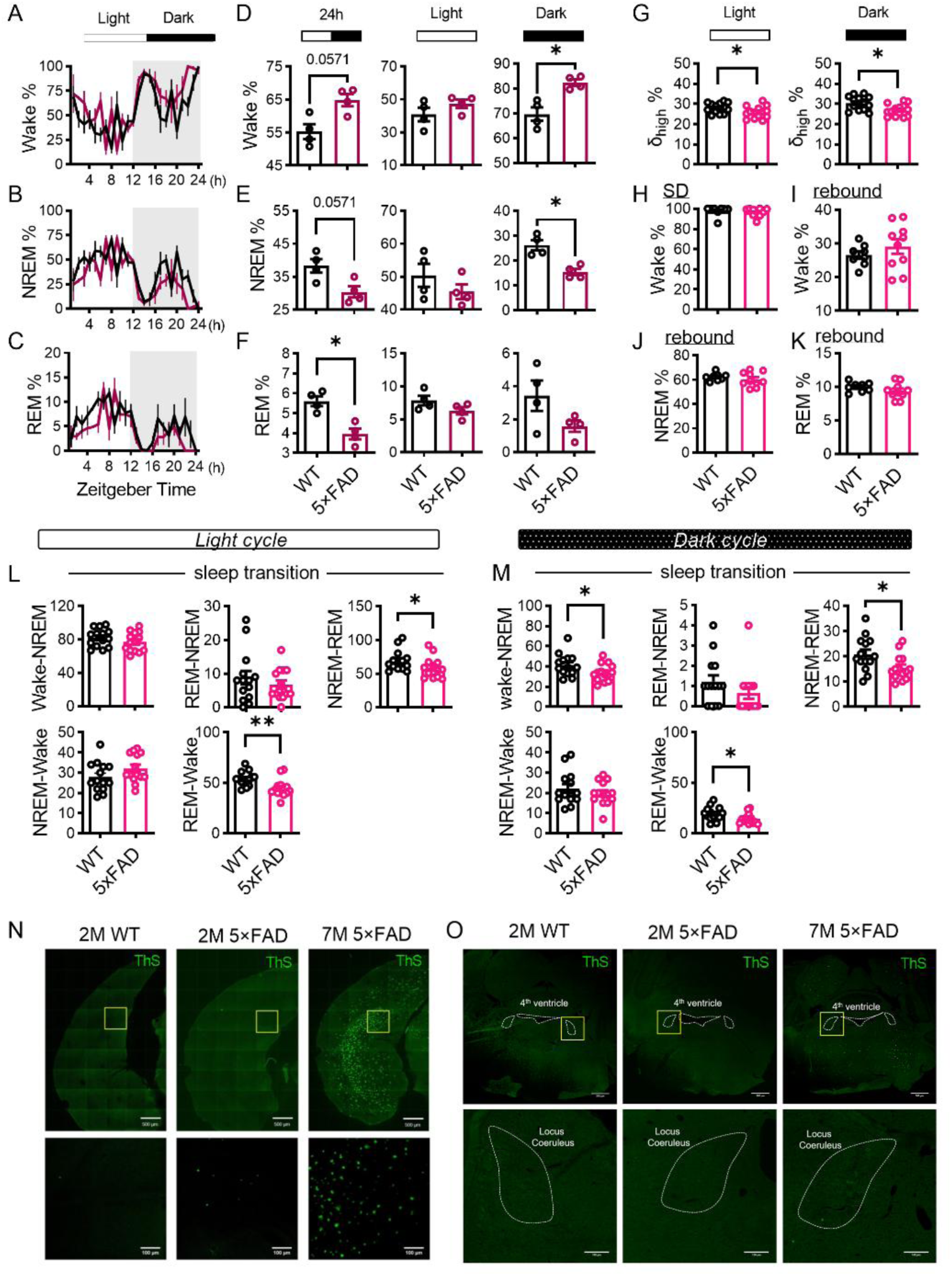
Sleep deficits in old and young 5xFAD mice. A-F. Percentages of time spent in wake (A), NREM (B), and REM (C) across the light/dark cycle in 12-month-old 5xFAD mice and their WT littermates. Comparisons of the percentage of time spent in wake (D), NREM (E), and REM (F) during the 24-hour day (left panels), light cycle (middle panels) and the dark cycle (right panels) between 12-month-old 5xFAD mice and their WT littermates. (WT: n = 4; 5xFAD: n = 4) G. Comparison of the percentages of δ_high_ (2-4 Hz) power of NREM sleep during the light cycle (left panel) and the dark cycle (right panel) between 2-month-old 5xFAD mice and WT mice. H-K. The 2-month-old 5xFAD mice exhibited normal sleep homeostasis. H, Wake percentages during the 4-hour sleep deprivation (SD) of the 2-month-old 5xFAD mice and WT mice. I-K, Wake (I), NREM (J) and REM (K) percentages during the 8-hour recovered sleep in 2-month-old 5xFAD and WT mice. (WT: n = 8; 5xFAD: n = 10) L-M. Changes in transition number between different vigilance stages during light cycle (L) and the dark cycle (M) in 2-month-old 5×FAD mice and WT controls. (WT: n = 14; 5xFAD: n = 14) N-O. Thioflavin S staining of amyloid plaques in hippocampus (N) and LC (O) of the 5xFAD mice of 2-month and 7-month of ages. (Yellow box in N: dentate gyrus; Yellow box in O: LC; Top panels scale bar: 500 µm; Bottom panels scale bar: 100 µm, dotted lines showing LC region)

**Figure S2.**
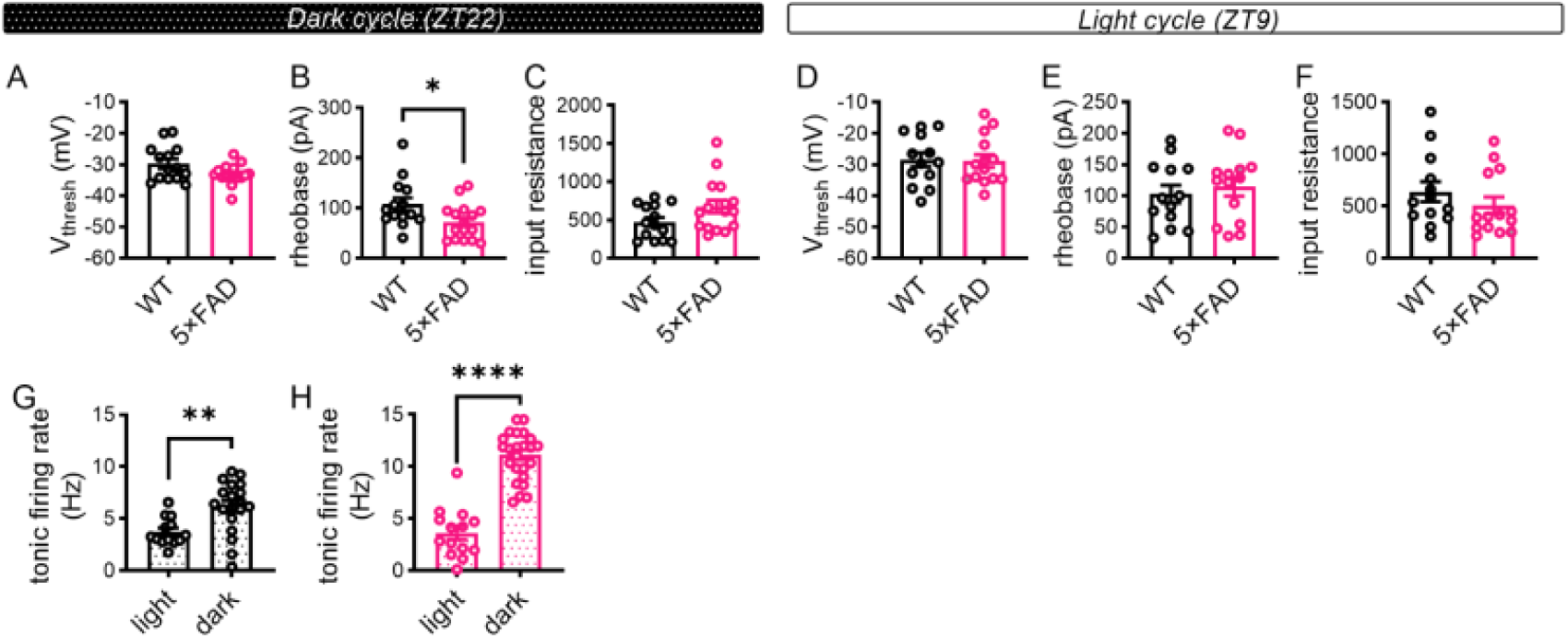
Electrophysiological properties of LC neurons in 2-month-old 5xFAD mice. A-C. Quantifications of action potential threshold potential (A), rheobase (B) and input resistance (C) of LC neurons in 2-month-old 5xFAD and WT mice during dark cycle. (WT: 14 cells from 3 mice; 5xFAD: 16 cells from 3 mice.) D-F. Quantifications of action potential threshold potential (D), rheobase (E) and input resistance (F) of LC neurons in 2-month-old 5xFAD and WT mice during light cycle. (WT: 13 cells from 3 mice; 5xFAD: 14 cells from 3 mice.) G-H. Comparison of tonic firing rate between light and dark cycle in 2-month-old WT mice (G) and 5xFAD mice (H) LC neurons.

**Figure S3.**
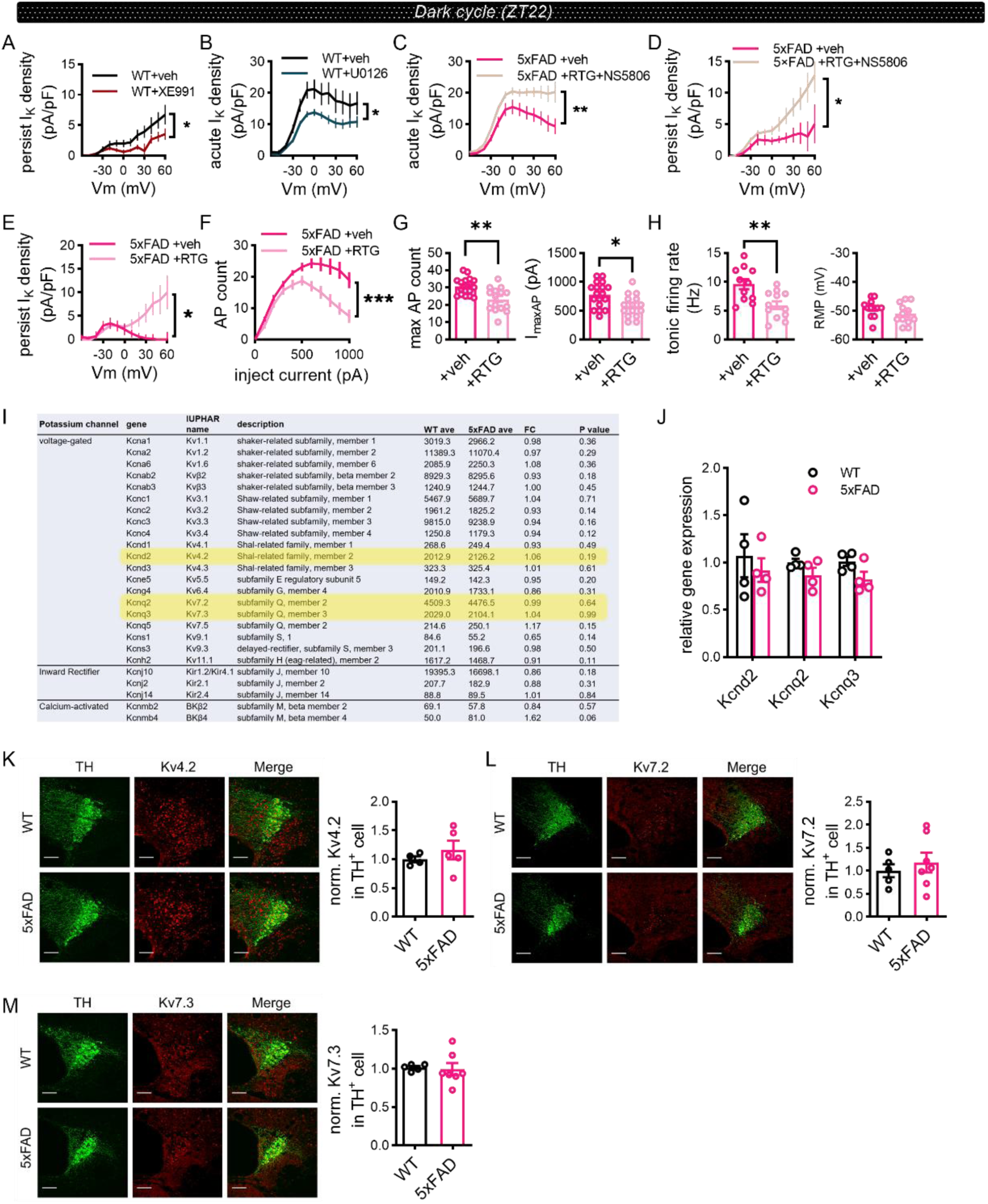
Impaired Kv4 and Kv7 channel function but not mRNA or protein expressions in young 5xFAD mice. A-D. The effect of XE991 (A, Veh: 10 cells from 3 mice; XE991: 13 cells from 3 mice.) or U0126 (B, Veh: 6 cells from 2 mice; U0126: 18 cells from 2 mice) on WT mice LC neurons and the effect of co-application of retigabine and NS5806 (C-D, Veh: 9 cells from 3 mice; RTG + NS5806: 13 cells from 3 mice.) on 5xFAD LC neurons low threshold I_k_. E-H. Effect of application of retigabine (RTG) on LT persist I_k_ (E), neuronal excitability (F-G) and tonic firing (H) of LC neurons of 5xFAD mice. E. low threshold persist Ik I-V curve. F. current-spiking relationship. G, left, maximal firing rate, right, I_maxAP_. H. Quantification of firing rate (left panel) and resting membrane potential (RMP, right panel). (E. Veh: 9 cells from 3 mice; RTG: 14 cells from 3 mice; F-G. Veh: 9 cells from 3 mice; RTG: 14 cells from 3 mice; H. Veh: 9 cells from 3 mice; RTG: 14 cells from 3 mice). I. Bulk sequencing of gene expression in the LC region of 2-month-old 5xFAD and WT mice revealed unchanged mRNA levels of a variety of potassium channels between 5xFAD and WT. (WT: n = 3 mice; 5xFAD: n = 3 mice) J, The relative gene expression levels of Kcnd2, Kcnq2 and Kcnq3 between 5xFAD and WT mice LC region (WT: n = 4 mice; 5xFAD: n = 4 mice) K-M. Immunofluorescence images showing Kv4.2 (K) Kv7.2 (L) and Kv7.3 (M) expression in the LC neuron of 5xFAD and WT mice (red: Kv4.2/ Kv7.2/ Kv7.3, green: TH, scale bar, 100 µm). Left panel, Representative immunofluorescence image; Right panel, Quantifications of normalized Kv channel signal intensity in TH positive cell. (K. WT: n = 4 mice; 5xFAD: n = 5 mice; L-M. WT: n = 5 mice; 5xFAD: n = 7 mice)

**Figure S4.**
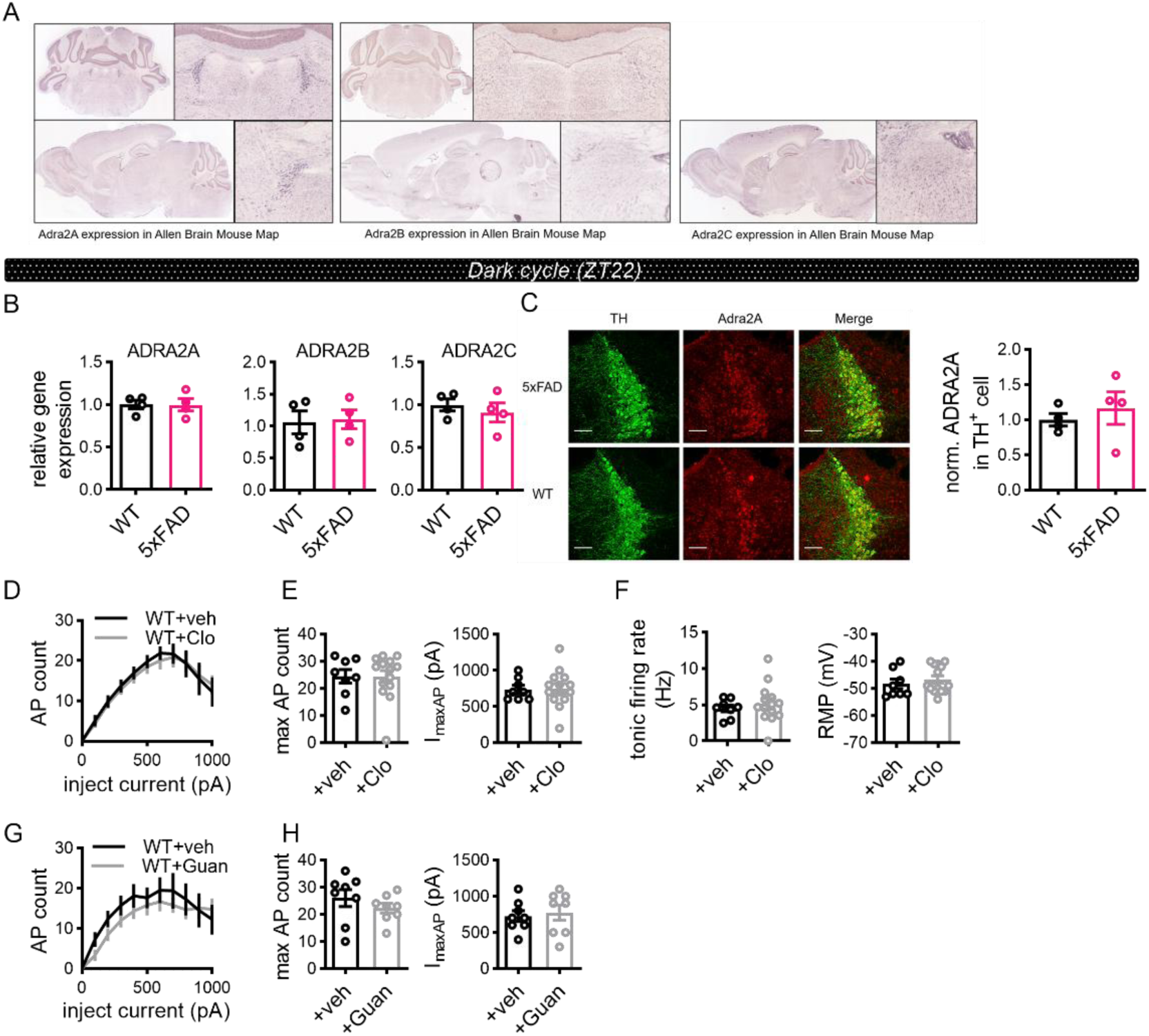
Unchanged mRNA or protein expression of α2A-AR suggest functionally impairment. A. Sagittal and coronal mouse brain atlas In Situ Hybridization (ISH) data showing ADRA2A, ADRA2B, ADRA2C expression in LC region from Allen Brain Atlas B. The relative gene expression levels of ADRA2A, ADRA2B and ADRA2C between 5xFAD and WT mice LC region. (WT: n = 4 mice; 5xFAD: n = 4 mice) C. Immunofluorescence images showing Adra2A expression in the LC of 5xFAD and WT mice (red: Adra2A, green: TH, scale bar, 100 µm). Left panel, Representative immunofluorescence image; Right panel, Quantifications of normalized Adra2A signal intensity in TH positive cell. (WT: n = 4 mice; 5xFAD: n = 4 mice) D-F. Effect of application of clonidine (Clo) on neuronal excitability (D-E) and tonic firing (F) of LC neurons of WT mice. D, current-spiking relationship. E, Left panel, maximal firing rate. Right panel, I_maxAP_ F, Left panel, tonic firing rate. Right panel, resting membrane potential. (Veh: 8 cells from 3 mice; Clo: 14 cells from 3 mice) G-H. Effect of application of guanfacine (Guan) on neuronal excitability of LC neurons of WT mice. G, current-spiking relationship. H, Left panel, maximal firing rate. Right panel, I_maxAP_ (Veh: 8 cells from 3 mice; Guan: 8 cells from 3 mice)

**Figure S5.**
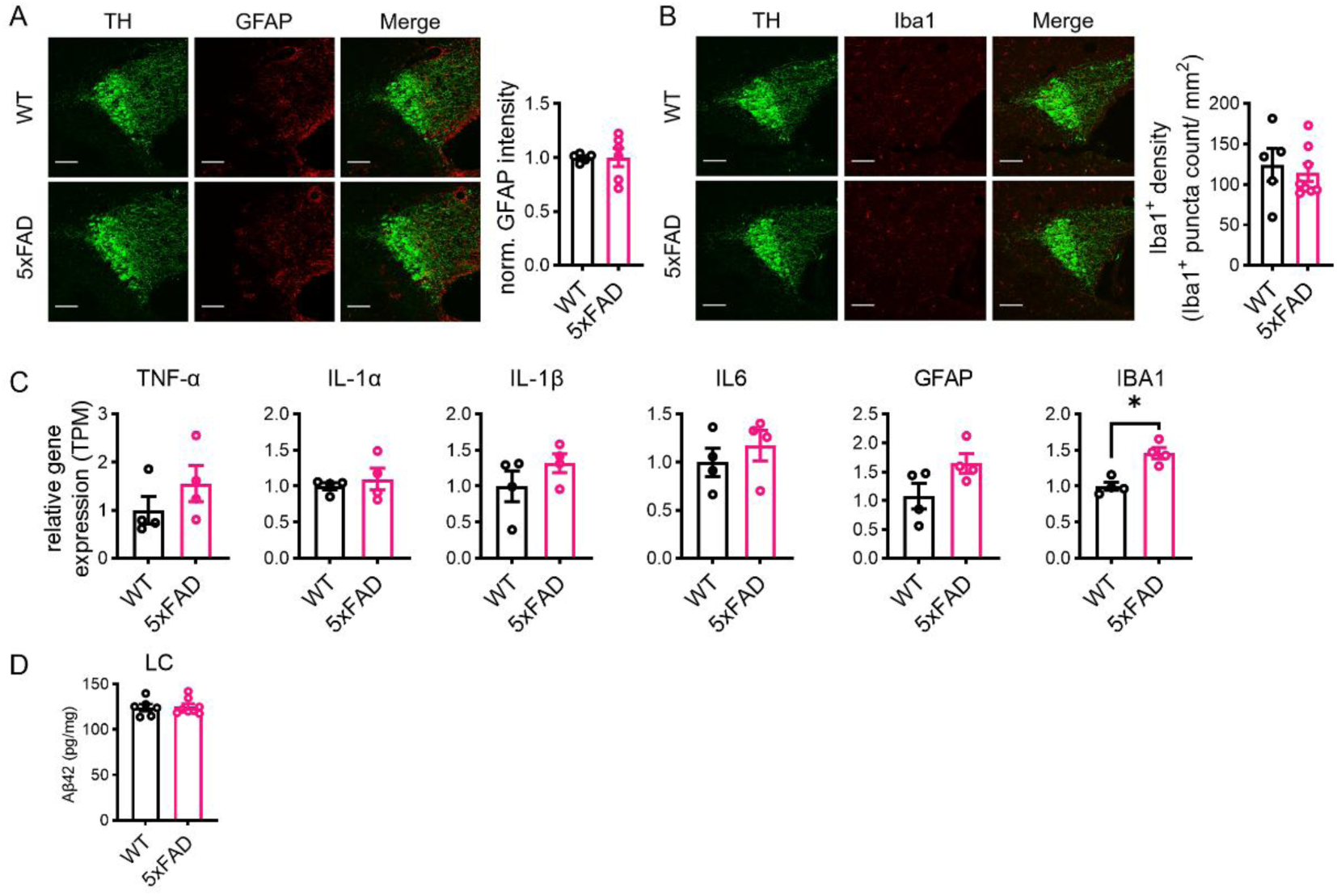
Mild neuroinflammation in LC region of 2-month-old 5xFAD mice. A-B. Immunolabeling GFAP (A) and Iba1 (B) of the LC neurons of 5xFAD and WT mice. Left, representative images (red: GFAP in A, Iba1 in B, green: TH, scale bar, 100 µm.). Right, comparisons of normalized GFAP signal intensity and Iba1+ density between 5xFAD and WT mice. (A, WT: n = 5 mice, 5xFAD: n = 6 mice; B, WT: n = 5 mice, 5xFAD: n = 8 mice) C. The relative gene expression levels of TNF-α, IL-1α, IL-1β, IL6, GFAP and IBA1 between 5xFAD and WT mice LC region (WT: n = 4 mice; 5xFAD: n = 4 mice) D. Comparison of Aβ42 in LC region between 5xFAD and WT. (WT: n=6 mice; 5xFAD: n=7 mice)

**Figure S6.**
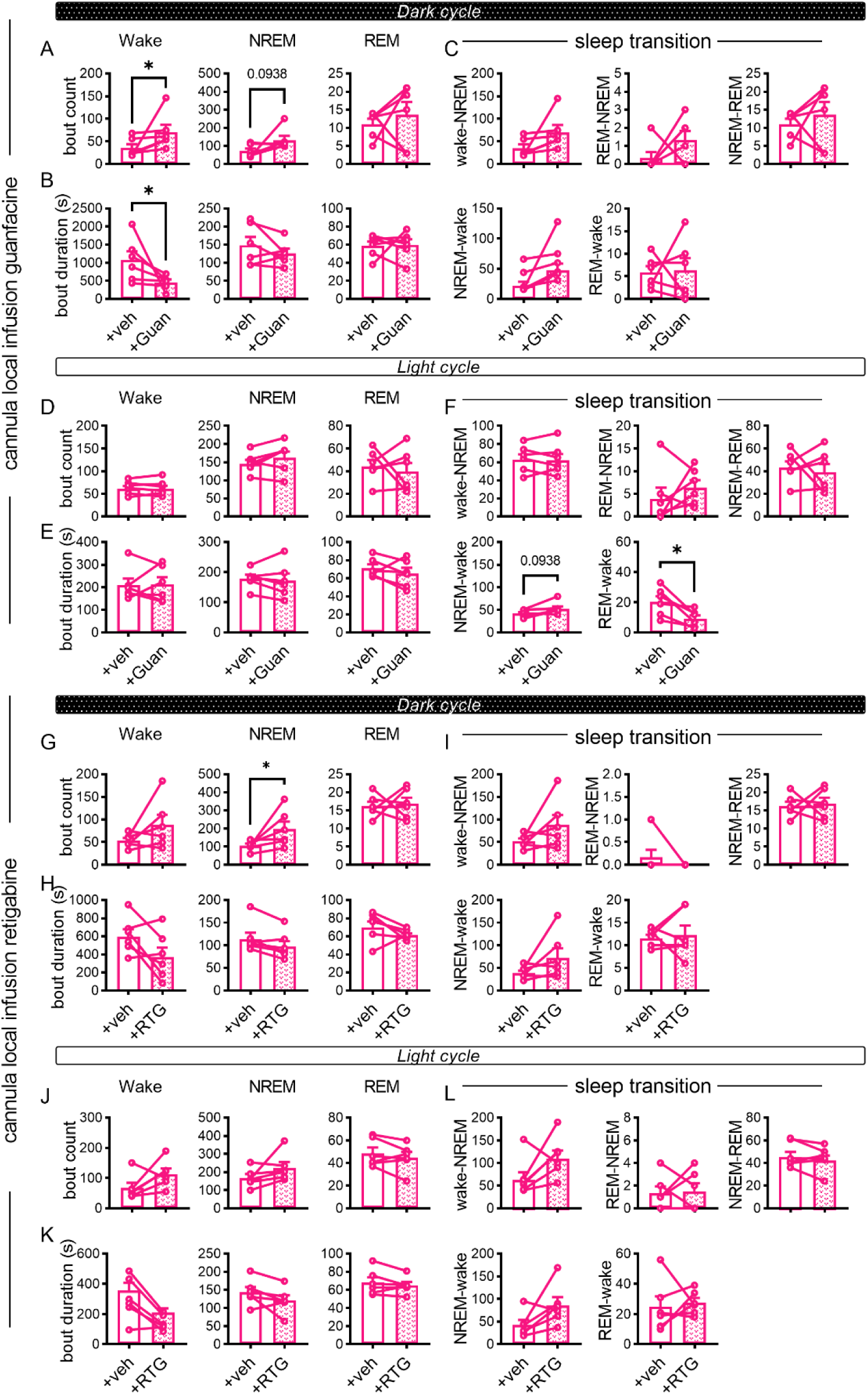
Local infusion of guanfacine or retigabine alleviates the sleep disturbances in 2-month-old 5xFAD mice. A-F. The differences in bout count number (A, D) and mean bout duration (B, E) for distinct vigilance stages and changes in transition number between different vigilance stages (C, F) during dark cycle (A-C) or the light cycle (D-F) in vehicle infusion (veh, opened bars) or guanfacine infusion (Guan, patterned bars) in 2-month-old 5×FAD mice with cannula implant in LC. A,B,D,E. Left panels, wake. Middle panels, NREM. Right panels, REM. (5xFAD: n = 6) G-L. The differences in bout count number (G, J) and mean bout duration (H, K) for distinct vigilance stages and changes in transition number between different vigilance stages (I, L) during dark cycle (G-I) or the light cycle (J-L) in vehicle infusion (veh, opened bars) or retigabine infusion (RTG, patterned bars) in 2-month-old 5×FAD mice with cannula implant in LC. G,H,J,K. Left panels, wake. Middle panels, NREM. Right panels, REM. (5xFAD: n = 6)

**Figure S7.**
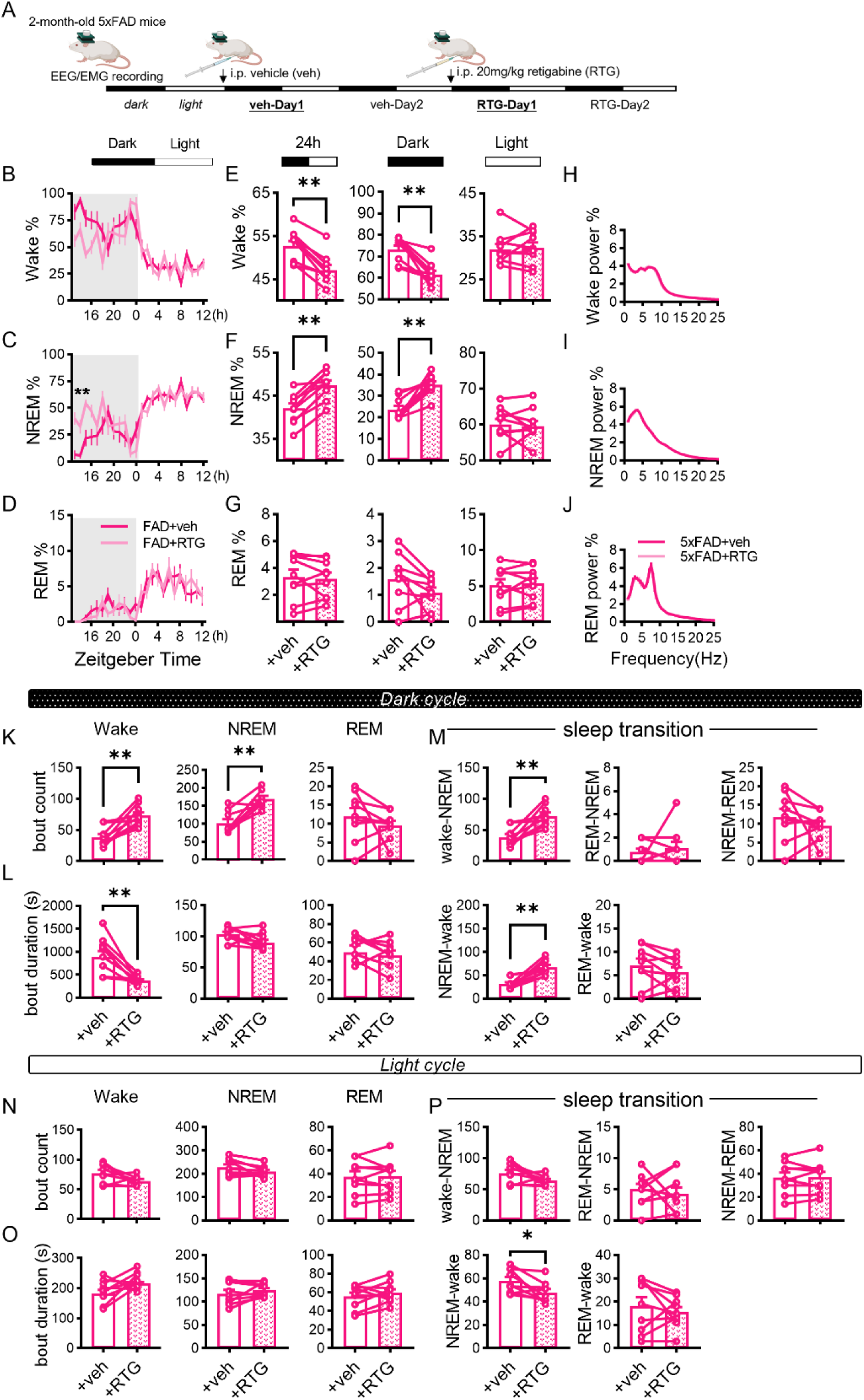
Systematic application of retigabine alleviates the sleep disturbances in 2-month-old 5xFAD mice. A. Schematic diagram showing the experimental design of intraperitoneal injection of retigabine (20 mg/kg) to evaluate effect on sleep. B-G. Comparisons of the percentages of time spent in wake (upper panels), NREM (middle panels), and REM (lower panels) immediately after i.p. of vehicle (veh, opened bars) and retigabine (RTG, patterned bars) in 5xFAD mice. B-D, Percentages of time spent in wake (B), NREM (C), and REM (D) across the light/dark cycle. E-G, Percentages in the 24-hour day (left panels), the dark cycle (middle panels) and the light cycle (right panels) were analyzed. (5xFAD: n=9) H-J. Comparison of the normalized EEG power spectrums of wake (H), NREM (I), and REM (J) during the 24-hour day between vehicle or retigabine treated 2-month-old 5xFAD mice. (5xFAD: n=9) K-P. The differences in bout count number (K, N) and mean bout duration (L, O) for distinct vigilance stages and changes in transition number between different vigilance stages (M, P) during dark cycle (K-M) or the light cycle (N-P) in vehicle i.p. (veh, opened bars) or retigabine i.p. (RTG, patterned bars) 2-month-old 5×FAD mice. K,L,N,O. Left panels, wake. Middle panels, NREM. Right panels, REM. (5xFAD: n = 9)

**Figure S8.**
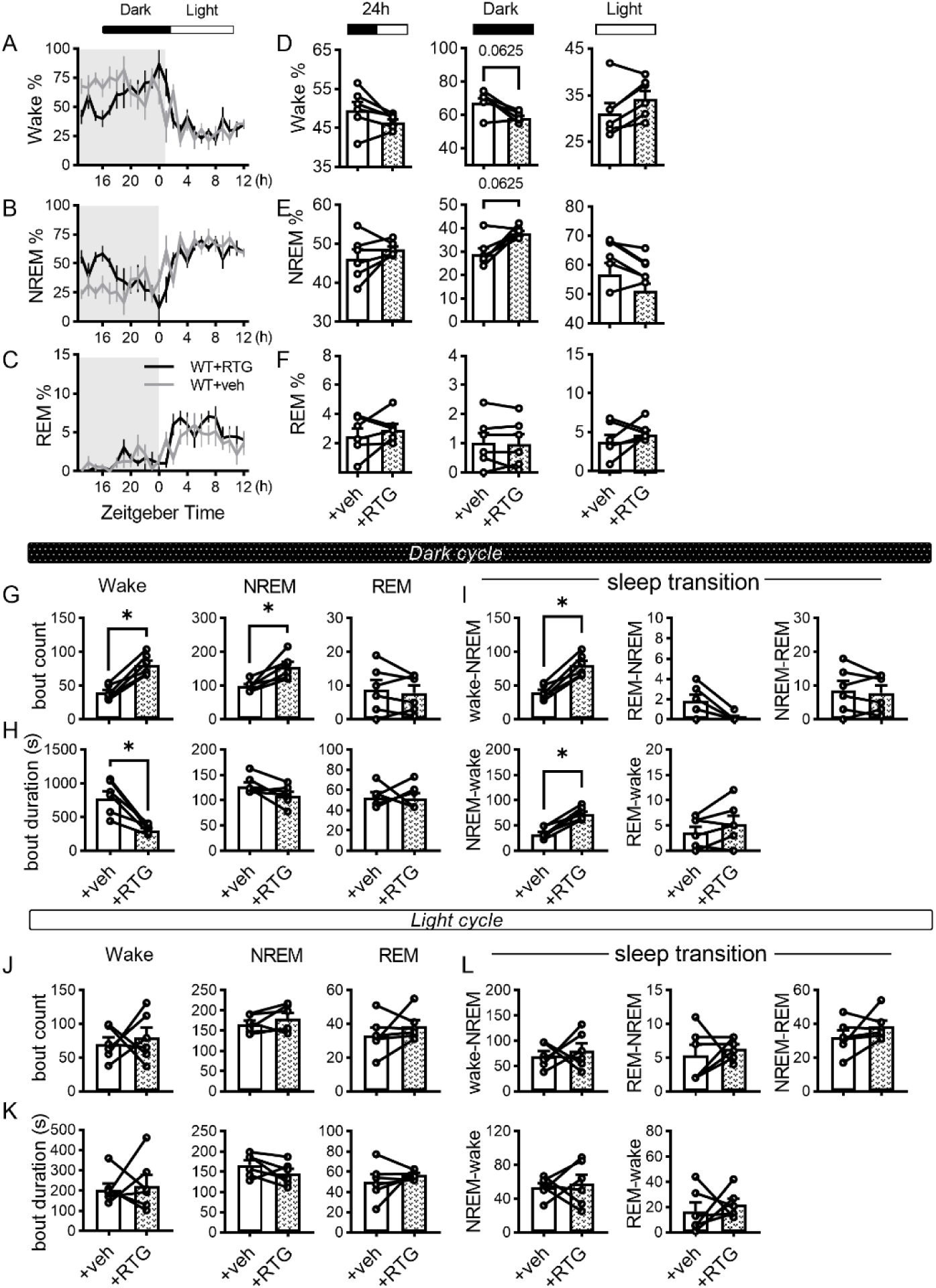
Systematic application of retigabine has mild sleep effect in 2-month-old WT mice. A-F. Comparisons of the percentages of time spent in wake (upper panels), NREM (middle panels), and REM (lower panels) immediately after i.p. of vehicle (veh, opened bars) and retigabine (RTG, patterned bars) in WT mice. A-C, Percentages of time spent in wake (A), NREM (B), and REM (C) across the light/dark cycle. D-F, Percentages in the 24-hour day (left panels), the dark cycle (middle panels) and the light cycle (right panels) were analyzed. (WT: n=6) G-L. The differences in bout count number (G, J) and mean bout duration (H, K) for distinct vigilance stages and changes in transition number between different vigilance stages (I, L) during dark cycle (G-I) or the light cycle (J-L) in vehicle i.p. (veh, opened bars) or retigabine i.p. (RTG, patterned bars) 2-month-old WT mice. G,H,J,K. Left panels, wake. Middle panels, NREM. Right panels, REM. (WT: n = 6)

## METHODS

### Animals

All procedures were approved by the Institutional Animal Care and Use Committees at the Interdisciplinary Research Center on Biology and Chemistry, Chinese Academy of Science. Mice were group-housed under a 12 h:12 h light/dark cycle with *ad libitum* access to food and water. Male 5xFAD mice (B6.Cg-Tg(APPSwFlLon,PSEN1*M146L*L286V)6799Vas/Mmjax, originally from Jackson Laboratory) with their wildtype littermates (referred to as WT) were utilized in this study. For all experiments except those in Fig S1A-F (12 month) and Fig S1N-O (7 month), mice of 2-3 months of age were used.

### Polysomnography recording

Polysomnography recording was carried out as described previously^67^. Briefly, mice were anesthetized by isoflurane vapor (1%–2%) and head-fixed. The skull was exposed and two epidural screws were implanted (B: +1.5 to -3 mm, L: +1.5 mm). Two insulated stainless-steel wires bared at the tip region were implanted into the dorsal right and left neck muscles and sutured in place to record the electromyogram (EMG). All electrodes were connected to a 4-pin socket connector that was cemented to the skull. Skin was sutured back and the wound was treated with triple antibiotic ointment and the mice were allowed to recover in their home cage for at least 10 days prior to recordings.

During recording, mice were transferred to the customized chamber with a day before the recording for habituation. EEG and EMG signals were recorded continuously depending on experiments. Regular 12 h:12 h light/dark cycle was maintained with *ad libitum* access to food and water. EEG and EMG signals were sampled at 256 Hz with 1000 times preamplifier gain and bandpass filtered at EEG: 0.3-1000 Hz, EMG: 1-5000 Hz. All signals were acquired by Sirenia Acquisition NiDAQ 1.7.9 (Pinnacle Technology).

For sleep deprivation, mice were kept awake by gentle handling during the first 4 hours (ZT0– 4) and allowed to sleep *ab libitum* during the remaining 8 h of that light cycle. EEG/EMG signals were acquired continuously.

To evaluate the effect of retigabine on sleep, 20 mg/kg of retigabine was administered by intraperitoneal (IP) injection approximately 30min before the onset of ZT12 followed by a complete 24-hour EEG/EMG recording. Same volume of vehicle was IP injected and the following EEG/EMG data from the same mouse was used as control.

### Cannula Implant

To monitor sleep with LC local drug application, bilateral cannula targeting LC area together with EEG/EMG electrode were implanted. Bilateral cannula was placed 200 μm above LC with following coordinations: Bregma, -5.45 mm; Lateral, ± 0.88 mm; Depth, -2.6 mm. All mice returned to their homecages post-surgery and were monitored every day for a week. 4µg retigabine (2 µg/µl) or 5.6 ng guanfacine (2.8 ng/µl) was infused into LC of each hemisphere approximately 30 min -1 hour prior ZT12 slowly via a micro-syringe (Hamilton).

### Slice Electrophysiology

#### Acute Brain Slices Preparation

Acute brain slices for electrophysiology were prepared as described previously ^68, 69^ 1-2 hours before the onset of the light cycle (designated as ZT22). Briefly, all mice were anesthetized with isoflurane vapor and transcardially perfused with about 10 ml of ice-cold dissection buffer containing the following: 93 mM NMDG, 2.5 mM KCl, 1.2 mM NaH_2_PO_4_·2H_2_O, 30 mM NaHCO_3_, 20 mM HEPES, 5 mM sodium ascorbate, 2 mM Thiourea, 3 mM sodium pyruvate, 25 mM D-Glucose, 12 mM NAc, 10 mM MgCl_2_, 0.5 mM CaCl_2_ bubbled with 95% O_2_/ 5% CO_2_. At the end of perfusion, mice were rapidly decapitated and whole brain was removed. 300 μm-thick acute coronal brain slices containing the Locus Coeruleus were sectioned by a vibratome (1200S, Leica) in ice-cold dissection buffer. Slices were incubated at in dissection buffer preheated to 30 °C for 15 min and then transferred to artificial cerebrospinal fluid (ACSF) containing: 119 mM NaCl, 5 mM KCl, 1.25 mM NaH_2_PO_4_, 1 mM MgCl_2_, 2 mM CaCl_2_, 26 mM NaHCO_3_, and 10 mM Dextrose, saturated with 95% O_2_ and 5% CO_2_). Slices were then allowed to recover in ACSF at room temperature for at least 1 h before recording.

#### Whole Cell Recording

For whole cell recordings, slices were transferred to a recording chamber continuously perfused with 30 ± 0.5°C ACSF at 2 ml/min. LC neurons were visualized by an upright fluorescence microscope (XT640-W, Olympus) and identified based on their localization and morphology. Borosilicate glass pipette recording electrodes with 3-6 MΩ tip resistance were filled with K^+^-based internal solution (130 mM K-gluconate, 10 mM KCl, 10 mM HEPES, 0.2 mM EGTA, 0.5 mM Na_3_GTP, 4 mM MgATP, 10 mM Na-phosphocreatine) adjusted to pH 7.2-7.4, 285-300 mOsm. Cells with a membrane resistance (Rm) > 100 MΩ and access resistance (Ra) < 25 MΩ were recorded. For all whole cell recordings, cells were discarded if these values changed more than 25% during the experiment. Data were filtered at 2 kHz and digitized at 10 kHz using Clampex (Axon).

### a. Spontaneous Firing

To measure spontaneous firing, cells were held at resting membrane potential with no current injection and spontaneous discharge was recorded continuously for at least 30 seconds. Only cells with resting membrane potential < -40 mV, Ra < 25 MΩ, and Rm > 100 MΩ were used for analysis.

### b. Spontaneous EPSCs and Neuronal Excitability

Recordings were done as previously described ^69, 70^. Both sEPSCs and excitability were recorded with regular ACSF without additional drugs. sEPSCs were recorded when cells were held at -60 mV in voltage clamp mode. To measure neuronal excitability, cells were current-clamped at -65 mV. Action potential threshold membrane potential and rheobase were estimated by injecting 500 ms ramp current from 0-400 pA. Maximal firing rate was measured by inject 500 ms current steps (0 to 1700 pA with 100 pA as step size). Only cells with resting membrane potential < -45 mV, Ra < 25 MΩ, and Rm > 100 MΩ were used.

### c. Miniature IPSCs

mIPSCs were recorded when cells were held at -70mV in voltage clamp mode with ACSF containing 1 μM TTX, 100 μM DL-APV and 20 μM CNQX. Recording electrodes were filled with mIPSC internal solution (130 mM CsCl, 8 mM NaCl, 0.2 mM CaCl2, 10 mM HEPES, 2 mM EGTA, 0.5 mM Na_3_GTP, 4 mM MgATP, 5 mM QX-314) adjusted to pH 7.2-7.3 with CsOH, 285-300 mOsm.

### d. Voltage-Gated Potassium Conductance

LC neurons were recorded in voltage clamp mode with K^+^-based internal solution. The following drugs were added to the perfused ACSF to isolate Kv currents: 1 μM TTX, 100 μM DL-APV, 20 μM gabazine and 20 μM CNQX. Total Kv currents were recorded by pre-holding cells at -120 mV for 750 ms followed by 400 ms step depolarization from -100 mV to +90 mV with 10 mV as step size. High threshold Kv currents were estimated by pre-holding cells at -30 mV for 750 ms prior to depolarization steps (Fig 3A). Low threshold Kv currents were calculated post hoc by subtracting high threshold Kv current from the total Kv current. Acute Kv currents were the maximal amplitude within 15 ms after current onset, whereas persist Kv currents were measured as the plateau amplitude at the end of each current step. For comparison, Kv current density was calculated by normalizing I_k_ to the membrane capacitance (Cm) of individual cell.

### e. Drug Application

Perfusion buffer containing 20 μM XE991, retigabine, U0126 or 10 μM NS5806 were used to evaluate the contribution of Kv7 or Kv4 channels to the Kv currents, excitability and spontaneous firing. 10 μM guanfacine and clonidine or 300nM BRL-44408 maleate were used to evaluate the contribution of α2A adrenergic receptor to the Kv currents, excitability and spontaneous firing.

#### Total Aβ measurement

Mice were anesthetized with isoflurane vapor and rapidly sacrificed. Brains were immediately removed. Brain tissues containing target regions were micro-dissected and weighted. Enzyme-linked immunosorbent assay (ELISA) kit specific for Aβ42 (#KHB3441, Invitrogen) was used to quantify total Aβ42 according to the manufacturer’s instructions.

#### Immunofluorescence Staining and Imaging Brain tissue preparation

Mice were anesthetized with isoflurane vapor and transcardially perfused with 5 ml PBS followed by 10 ml of 4% phosphate buffered paraformaldehyde (PFA) for immunofluorescence. Brains were then removed and fixed overnight in PFA.

#### Immunofluorescence staining (IF)

PFA-fixed brains were dehydrated in 20% phosphate-buffered sucrose for 1 day and 30% phosphate-buffered sucrose for 2 days before embedded in tissue freezing medium (#14020108926, Leica) and rapidly frozen in liquid nitrogen. 20 μm-thick coronal brain slides containing LC or hippocampus were section by Cryostat (CM3050 S, Leica). For staining, the slices were first treated with frozen slice antigen retrieval reagent (#P0090, Beyotime). They were then blocked with PBS containing: 3% NDS (Normal Donkey Serum), 0.3% Triton X-100 at room temperature for 1 hour before incubated with primary antibodies (TH, 1:3000; NeuN, 1:2000; Kv4.2, 1:500 Kv7.2, 1:500; Kv7.3, 1:500; GFAP, 1:10000; Iba1, 1:1000, Adra2A, 1:1000) overnight at 4°C. After washing with PBST, slices were incubated with secondary antibodies at room temperature for 2 hours. Slices were washed with PBST and then mounted on glass slide and let air dry. Prolong Gold Antifade Mountant (#P36930, Invitrogen) was used to slow down bleaching of fluorescence. Amyloid plaque was stained by using Thioflavin S in combined with a gradient ethanol treatment following the manual instruction.

All antibodies and materials used are listed in Table S1.

#### Confocal Imaging

All fluorescence signal was acquired using a spinning disk microscope (Dragonfly 200, Andor) connected to camera (Zyla sCMOS, Andor) with a 20x air lens. The image resolution is x/y: 0.300903/ 0.301040 μm/pixel. Stack images were acquired for all images. Step size (μm) and thickness (μm) of were as followed: 0.5, 1

#### Tissue Preparation, RNA extraction and Bulk RNA sequencing

Mice were anesthetized with isoflurane vapor and rapidly sacrificed. Brain slices containing LC region were collected by a vibratome (1200S, Leica) in ice-cold dissection buffer. LC regions were micro-dissected on ice with sterile disposable biopsy punch (#33-31, Integra Lifesciences) then quickly frozen with liquid nitrogen and stored at −80°C until used. TRIzol^®^ Reagent (Invitrogen Life Technologies) was used to retrieve total RNA. RNA concentration and quality were determined by NanoDrop spectrophotometer (Thermo Scientific). Only high-quality RNA samples (OD260 / 280 = 1.8 ∼ 2.2, OD260 / 230 ≥ 1.9, RIN ≥ 9.0) were used for the library and bulk RNA sequencing (Shanghai Personal Biotechnology Cp. Ltd.).

#### Quantitative real-time polymerase chain reaction (qPCR)

For qPCR, after DNase treatment to remove genomic DNA, the reverse transcription reactions were performed using Hifair III 1st Strand cDNA Synthesis SuperMix kit (#11141ES60, Yeason). The cDNA was then used for real-time qPCR using TB Green Premix Ex Taq II (#RR820A, Takara) with the QuantStudio 6 Flex Real-Time PCR system (Life Technologies). The qPCR data were calculated using the 2-ΔΔCt method. The mRNA levels of actin were used as an internal control to normalize the mRNA levels of genes of interest.

The following qPCR primers were used in this study: *mActin*-F, 5′-GTGTGATGGTGGG AATGGGT-3′; *mActin-*R, 5′-GCTGGGGTGTTGAAGGTCTC-3′; *mTNFα-*F: 5′-CCCTCACA CTCAGATCATCTTCT-3′; *mTNFα*-R: 5′-GCTACGACGTGGGCTACAG-3′; *mIL1α*-F, 5′-CG AAGACTACAGTTCTGCCATT-3′; *mIL1α-*R, 5′-GACGTTTCAGAGGTTCTCAGAG′; *mIL1β*-F, 5′-GCAACTGTTCCTGAACTCAACT -3′; *mIL1β-*R, 5′-ATCTTTTGGGGTCCGTCAACT -3′; mIL6-F, 5′-TAGTCCTTCCTACCCCAATTTCC -3′; *mIL6*-R, 5′-TTGGTCCTTAGCCACTCCTTC -3′; *mGfap-*F, 5′-CGGAGACGCATCACCTCTG-3′; *mGfap*-R, 5′-AGGGAGTGG AGGAGTCATTCG-3′; *mIba1*-F, 5′-ATCAACAAGCAATTCCTCGATGA-3′; *mIba1*-R, 5′-CA GCATTCGCTTCAAGGACATA -3′; *mKcnd2*-F, 5′-GGGTGGATGCCTGTTGCTT-3′; *mKcnd 2*-R, 5′-GTCTTGCCATGTCTGGAAACG-3′; *mKcnq2*-F, 5′-CTGCCTGGAGATTCTATGCT ACT -3′; *mKcnq2*-R, 5′-AGTGACTGTCCGCTCGTAGT-3′; *mKcnq3*-F, 5′-GAGCCGACAA AGACGGGAC -3′; *mKcnq2*-R, 5′-TTGGCGTTGTTCCTCTTGACT -3′; *mAdra2a*-F, 5′-G TGACACTGACGCTGGTTTG -3′; *mAdra2a*-R, 5′-CCAGTAACCCATAACCTCGTTG -3′; *m Adra2b*-F, 5′-TCTTCACCATTTTCGGCAATGC -3′; *mAdra2b*-R, 5′-AGAGTAGCCACTAG GATGTCG -3′; *mAdra2c*-F, 5′-GACGCAAGCGGTAGAGTACA -3′; *mAdra2c*-R, 5′-GTAG AACGAGACGAGAGGCG-3′

## QUANTIFICATION AND STATISTICAL ANALYSIS

### Analysis of sleep-wake states

States of wakefulness, NREM, REM sleep, and microarousal were first automatically analyzed in 5-second epoch by NeuroScore (Data Sciences International) using the same criteria: Slow Wave Ratio: 0.4, Theta Ratio: 3. The results were manually checked. The wake epochs shorter than 10 sec were defined as microarousal.

### EEG signal processing and analysis

EEG signals were band-pass filtered at 0.5-100 Hz and power line filtered at 50 Hz before analyzed by Fourier Transform. The brain oscillation bands were defined as Delta (δ): 0.5-4 Hz, Theta (θ): 4-8 Hz, Alpha (α): 8-12 Hz, Beta (β): 12-30 Hz, Gamma_low_ (γ_low_): 30-50 Hz. The Unitary power was defined as average power per 5-second epoch. Each bout is defined as a continuous state of wake/ NREM/ REM with no interruptions for sleep structure analysis.

### sEPSCs and mIPSCs analysis

sEPSCs and mIPSCs were analyzed using the MiniAnalysis program (Synaptosoft, Decatur, GA) as described previously ^69^. Event detection threshold was set at 3 times over the RMS noise. At least 300 events with rise time < 3 msec were selected for each cell to calculate sEPSC and mIPSCs frequency and amplitude.

### Image Analysis

Immunofluorescent images were batched analyzed by Fiji ImageJ (NIH) and Imaris (Oxford Instruments). All images were projected in z direction with max intensity followed by background subtraction. For intensity signals, LC neurons were identified by TH positive signals and the positive region signal were measured. For Iba1 positive puncta density measurements, Iba1 positive puncta were automatically selected and counted by Spot module of Imaris. All images were analyzed in a double-blinded manner. To report the immunofluorescence results, the mean signal intensity was normalized to control groups. For different batches of experiments, groups were normalized to their own control before combination. Raw value of Iba1 puncta density was reported.

### Bulk RNA sequencing analysis

The Bioinformatics pipeline (https://github.com/emc2cube/Bioinformatics) was used to process sequencing data. STAR^71^ was used to align reads to mm39 genome build and RSEM was used to quantify expression at the gene level^72^. Differential expression analysis was conducted using DESeq2 with default parameters^73^. Differential gene expression was identified using a statistical cut-off of padj < 0.05

### Statistics

Sample size is indicated in all figures as mouse number or cell then mouse number. Statistical analysis was performed by Prism V6.0 software (GraphPad Software, Inc.). Wilcoxon signed-rank test was used for paired data. Mann-Whitney U test was used for unpaired data. Two-way *ANOVA* with Sidak post hoc multiple-comparisons test were used to compare I-V curves and I-O curves. Error bars in all figures indicate standard error of mean (s.e.m). Significant comparisons were labeled in the figures. The level of significance was set at P < 0.05. *P < 0.05; **P < 0.01; ***P < 0.001; ****P<0.0001. P values (0.05<P<0.1) were labeled in graph.

### Code Availability

The customized MATLAB codes are available on GitHub. https://github.com/KaiWen-Helab/Zhang_tau2024

### KEY RESOURCES TABLE

**Supplementary Table 1.**
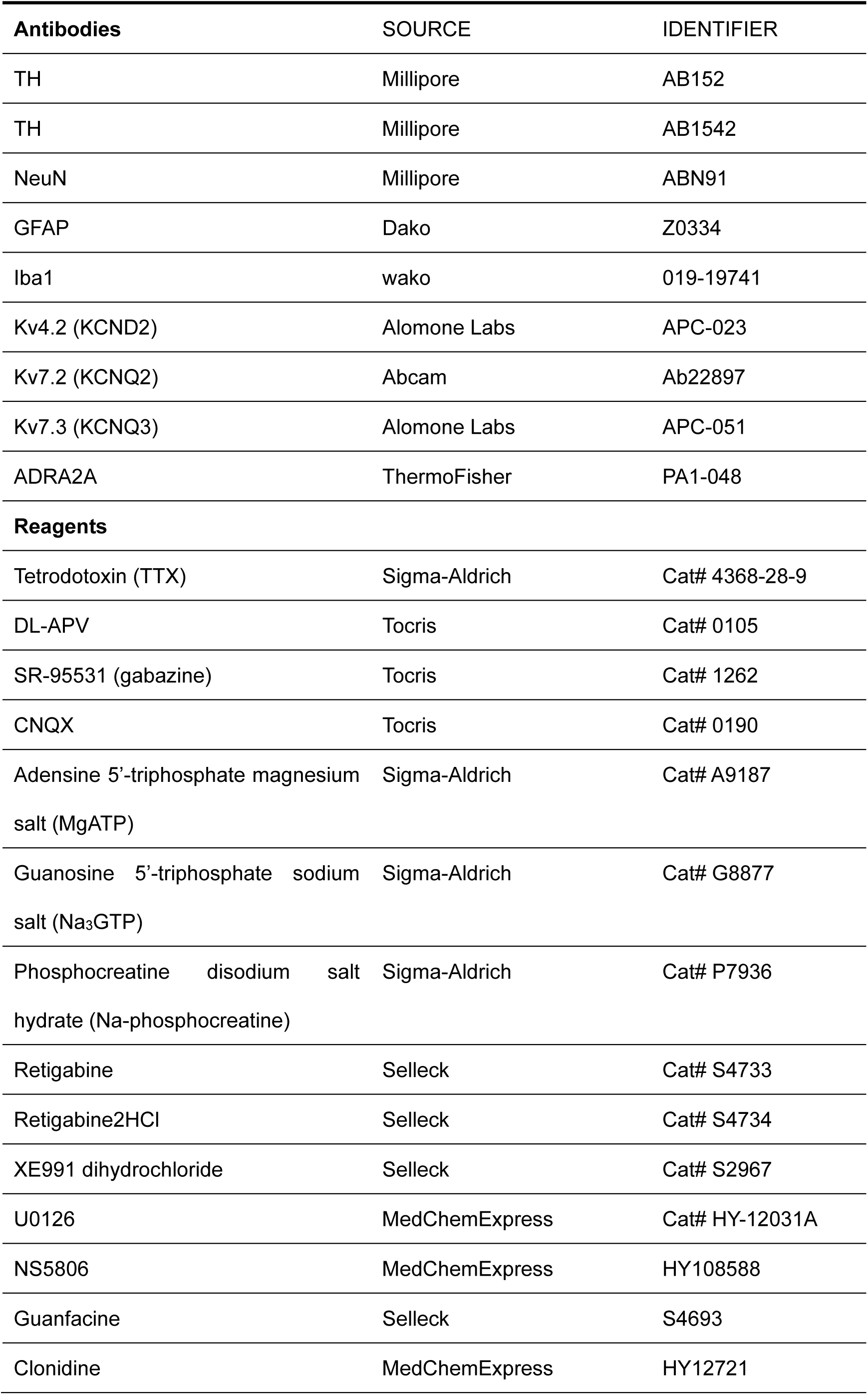
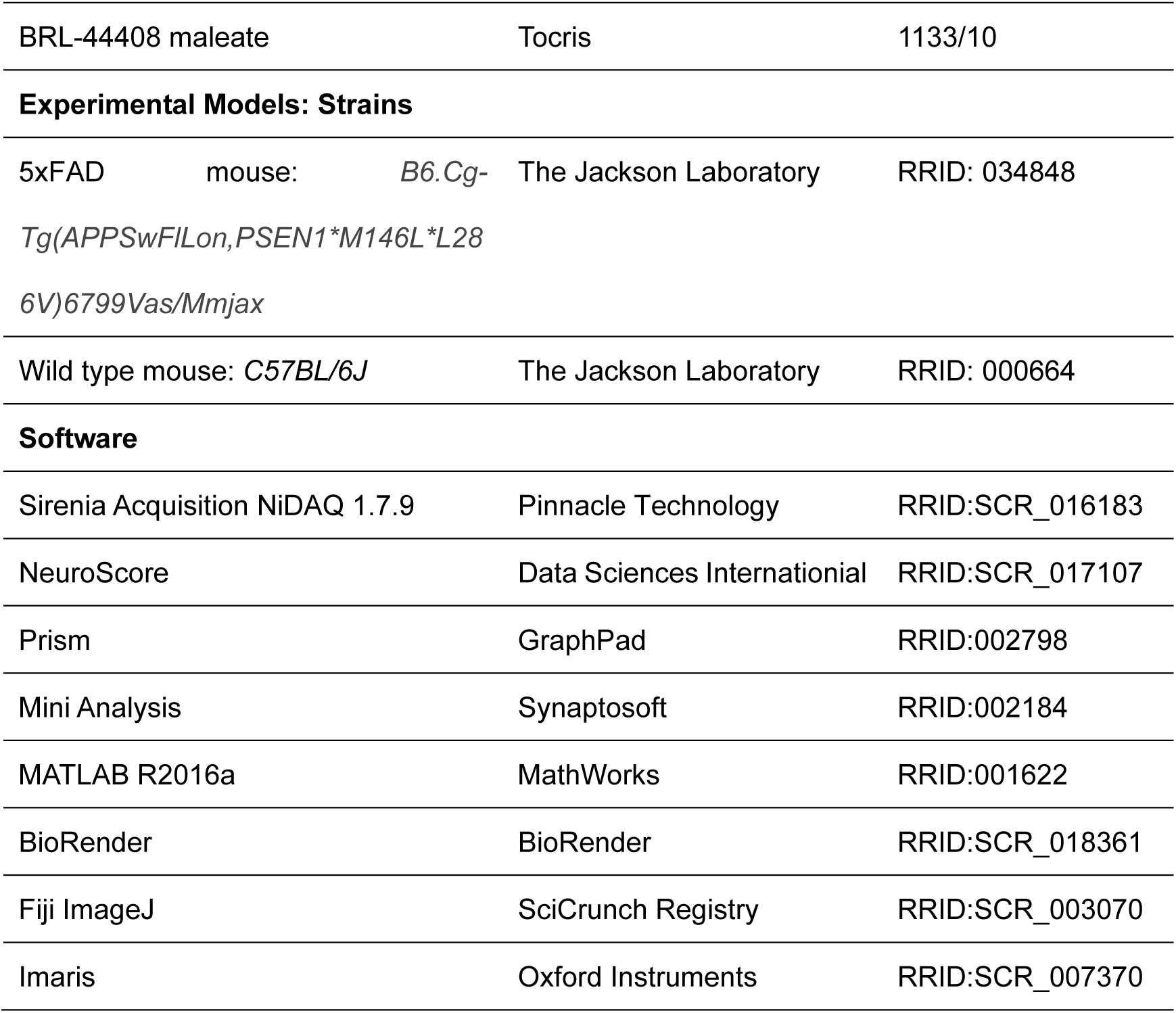
Key resources table.

